# Genetic ablation of GABA_B_ receptors from oligodendrocyte precursor cells protects against demyelination in the mouse spinal cord

**DOI:** 10.1101/2024.05.29.596385

**Authors:** D Gobbo, P Rieder, LP Fang, E Buttigieg, M Schablowski, E Damo, N Bosche, E Dallorto, P May, X Bai, F Kirchhoff, A Scheller

## Abstract

GABAergic signaling and GABA_B_ receptors play crucial roles in regulating the physiology of oligodendrocyte-lineage cells, including their proliferation, differentiation, and myelination. Therefore, they are promising targets for studying how spinal oligodendrocyte precursor cells (OPCs) respond to injuries and neurodegenerative diseases like multiple sclerosis. Taking advantage of the temporally controlled and cell-specific genetic removal of GABA_B_ receptors from OPCs, our investigation addresses their specific influence on OPC behavior in the gray and white matter of the mouse spinal cord. Our results show that while GABA_B_ receptors do not significantly alter OPC cell proliferation and differentiation under physiological conditions, they distinctly regulate the Ca^2+^ signaling of OPCs. In addition, we investigate the impact of OPC-GABA_B_ receptors in two models of toxic demyelination, namely the cuprizone and the lysolecithin models. The genetic removal of OPC-GABA_B_ receptors protects against demyelination and oligodendrocyte loss. Additionally, we observe enhanced resilience to cuprizone-induced pathological alterations in OPC Ca^2+^ signaling. Our results provide valuable insights into the potential therapeutic implications of manipulating GABA_B_ receptors in spinal cord OPCs and deepen our understanding of the interplay between GABAergic signaling and spinal cord OPCs, providing a basis for future research.

## Introduction

In the central nervous system (CNS), oligodendrocyte precursor cells (OPCs) represent a reservoir of new mature oligodendrocytes and exist in a 1:4 ratio with myelinating oligodendrocytes (*1–5*). OPCs rapidly proliferate in response to injury (*6*), such as acute spinal cord injury (*7*), or in the context of neurodegenerative diseases affecting spinal cord myelination including chronic inflammatory diseases such as multiple sclerosis (MS) (*8, 9*). The loss of myelin occurring during MS is associated with immune-driven oligodendrocytes death (*10–12*) and results in axon exposure and subsequent neurodegeneration (*13, 14*). As a response, OPCs migrate to the white matter lesions driven by resident micro- and astroglia, proliferate and differentiate (*15–18*) and participate in the spontaneous regeneration of myelin (*19, 20*).

In line with the ability of OPCs to form synapses with neurons, both glutamate and GABA (*21, 22*) have been identified as regulators of the oligodendrocyte-lineage cell proliferation (*23–26*), differentiation (*23, 24, 26*) and maturation (*25*) as well as myelination (*24, 25, 27, 28*). In particular, recent studies focusing on interneuron myelination and interneuron-OPC synapses in the brain have shed light on the importance of GABA signaling in the oligodendrocyte lineage (*28, 29*). The activation of OPC GABA_B_ receptors (GABA_B_Rs) increases OPC proliferation and migration *in vitro* (*26, 30*). Also, the GABA_B_R agonist baclofen enhances remyelination in the lysolecithin model of spinal cord demyelination *in vivo* (*31*). GABA_B_R antagonism increases OPC proliferation and decrease OPC maturation and myelination in the rat brain during development (*32*). In OPCs, activation of GABA_B_Rs results in G_i/o_ protein-induced reduction of intracellular levels of cyclic AMP (*30*) but an alternative G_q_ protein-pathway linked to phospholipase C and intracellular Ca^2+^has been suggested as well (*22*). Indeed, OPCs respond to GABAergic neurons via intracellular Ca^2+^ oscillations (*33, 34*). Also, Ca^2+^ signaling has been linked to the regulation of myelination in oligodendrocyte-lineage cells (*35–38*) as well of OPC fate (*39, 40*).

The involvement of GABA_B_Rs as well as OPC Ca^2+^ signaling in the regulation of oligodendrocyte-lineage cell proliferation and fate points at their contribution to the pathological outcome of demyelination insults. Indeed, both pre- and postsynaptic GABAergic neurotransmission (*41, 42*) as well as GABA levels (*43, 44*) are altered in the brains of MS patients and the GABA_B_R-agonist baclofen itself is currently used as a therapeutical agent for spasticity in MS (*45*). The specific role of GABA_B_Rs in OPCs within spinal cord physiology and pathology remains poorly understood *in vivo*. In this study, we address this gap by employing genetic removal of GABA_B_Rs from OPCs, aiming to elucidate their contribution to cell proliferation, differentiation, myelination, and OPC Ca^2+^ signaling. Additionally, taking advantage of the cuprizone and lysolecithin models of toxic demyelination, we explore the role of OPC GABA_B_Rs in the cellular response to pathological demyelinating insults of the spinal cord white matter.

## Materials and Methods

### Animals

Mice were maintained in the animal facility of the Center for Integrative Physiology and Molecular Medicine (CIPMM, University of Saarland, Homburg). Humidity and temperature were maintained at 45–65 % and 20–24°C and the facility was kept under a 12 h light-dark cycle. All mice received food *ad libitum* (standard autoclaved rodent diet, Ssniff Spezialdiäten, Soest, Germany) and autoclaved tap water. Knock-in NG2-Cre^ERT2^ mice (Cspg4^tm1.1(cre/ERT2)Fki^, MGI: 5566862) (*4*) were crossbred to mice with Rosa26 reporter mice (Gt(ROSA)26Sor^tm1(CAG-GCaMP3)Dbe^, MGI: 5659933) (*46*) and Gabbr1^tm2Bet^ mice (MGI: 3512742) (*47*) and maintained in a C57BL/6N background. To induce genetic recombination, tamoxifen was administered intraperitoneally (i.p.) for five consecutive days (once per day, 100 mg/kg body weight) (*48*) at 10 weeks of age. In Supplementary Figure S1, early tamoxifen administration was performed at one week of age (two consecutive days, once per day, 100 mg/kg body weight) or at four weeks of age (five consecutive days, once per day, 100 mg/kg body weight). Two-photon laser-scanning microscopy (2P-LSM) and immunohistochemistry (IHC) were performed at least two weeks after beginning of the tamoxifen treatment. For the demyelination study, 12-week-old male and female mice were fed with standard powder food containing 0.3 % cuprizone for one week and 0.2 % cuprizone for other two weeks (Bis(cyclohexanone)oxaldihydrazone, Cat. No. CC99924, Carbolution Chemicals, St. Ingbert, Germany). The diet was freshly prepared and changed every day. After three weeks of diet, the animals were fed standard food until further analysis. This protocol has been previously established and was used to address demyelination in the brain resulting in a 90 % loss of oligodendrocytes in the middle part of the corpus callosum (*49*). 2P-LSM and IHC analysis for cuprizone treated-mice and age matched controls were performed on 15-16 week old mice.

### Laminectomy and spinal window implantation

All surgical sections were realized in animals under inhalational anesthesia (1.5-2 % isoflurane, 66 % O_2_ and 33 % N_2_O) and the animaĺs eyes were covered by Bepanthen (Bayer, Leverkusen, Germany). Surgeries were executed as previously described (*50*) to get access to T12-L2 vertebrae and by laminectomy approach, L4-S1 spinal segments could be exposed. A modified coverslip was fit on the spinal cord and animals were treated with analgesic and antiphlogistic agents for three consecutive days (*51*).

### Acute lysolecithin incubation

The procedure for acute incubation with lysolecithin (LPC, 1 % v/v in NaCl 0.9 %) was performed as previously described (*52*). After laminectomy, the exposed *dura mater* was opened with a fine needle, washed with artificial cerebrospinal fluid (aCSF) and incubated for one hour with 20 μl LPC. Afterwards, the LPC was dried out and the tissue was thoughtfully washed with aCSF. The window was placed and sealed as previously described. Animals were perfused one week later.

### Two-photon laser-scanning microscopy

*In vivo* recordings were performed using a custom-made two-photon laser-scanning microscope (2P-LSM), equipped with a mode-locked Ti:sapphire femtosecond pulsed laser, Vision II (Coherent, St. Clara, USA) (*53*), in combination with ScanImage software (*54*) as previously described (*50*). For transgenic GCaMP3 excitation, the laser wavelength was set to 890 nm and the power was adjusted from 8 to 60 mW, depending on the imaging depth in the tissue. 2P-LSM was performed on the white matter of the dorsal funiculus up to a 100-150 µm depth by using a long-distance W Plan-Apochromat 20x 1.0 NA DIC objective (Zeiss, Oberkochen, Germany). Areas of white matter were recorded as uniformly spaced planes of field of views with 256×256 pixel per image, 1.4 µs pixel dwell time and GCaMP3 emission was acquired using a 500/24 nm band pass filter, detected by a photomultiplier tube H10770PB-40 (Hamamatsu Photonics, Hamamatsu, Japan). During 2P-LSM *in vivo* recordings, animals were kept under inhalation anesthesia with 1.5 % isoflurane on a heating plate.

### Automated ROA-based detection and analysis of Ca^2+^ events

Ca^2+^-event analysis was performed using the custom-made MATLAB-based analysis software MSparkles (*55*) as previously described (*50*). Shortly, fluorescence fluctuations at basal Ca^2+^ concentrations (F_0_) were computed along the temporal axes of each individual pixel, by fitting a polynomial of user-defined degree in a least-squares sense. The range projection of ΔF/F_0_ was then used to identify local fluorescence maxima, serving as seed points for simultaneous, correlation-based region growing (minimum ROA area, 5 µm^2^; temporal correlation threshold, 0.2). Prior to F_0_ estimation, image stacks were denoised using the PURE-LET algorithm (*56*) as well as a temporal median filter to correct small motion artefacts and simultaneously retain sharp transient edges. Based on the pre-processed data (F), Ca^2+^ event detection and analysis were performed on the normalized dataset (ΔF/F_0_).

### Immunohistochemistry

Anesthetized animals were transcardially perfused with phosphate-buffered saline (PBS) and tissue was fixed by 4 % formaldehyde perfusion. After 24 h post fixation in 4 % formaldehyde, T13-L1 spinal cord segments were dissected and detached from meninges. The spinal cord tissue was maintained in PBS and cut in transversal sections (40 µm) using a vibratome (VT1000 S) (Leica, Nußloch, Germany). Free floating slices were processed as described before (*57*). Briefly, incubation in blocking solution (Triton X-100, horse serum and PBS) at room temperature (RT) was followed by primary antibody solution incubation overnight at 4 °C for detection of the following markers: monoclonal mouse: anti-APC CC-1 (1:50, Cat. No. OP80, Calbiochem); anti-MBP (1:500, SMI99, Biolegend); monoclonal rat: anti-Ki67 (1:500, Cat. No. 14-5698-82, Thermo Fisher Scientific); polyclonal rabbit: anti-MyRF (1:500, Oasis Biofarm, Cat. No. OB-PRB007), anti-Olig2 (1:1000, Cat. No. AB9610, Millipore); polyclonal goat: anti-GFAP (1:1000, Abcam, ab53554), anti-Iba1 (1:1000, Abcam, ab5076), anti-PDGFRα (1:1000, Cat. No. AF1062, R&D Systems); polyclonal chicken: anti-GFP (1:1000, Cat. No. 10524234, Thermo Fisher Scientific), anti-NF (1:500, ab4680, Abcam). For MBP myelin staining, sections were pretreated with ethanol (> 99 %) for 10 min prior to incubation in blocking solution. Secondary antibodies (donkey secondary antibodies conjugated with Alexa488, Alexa555, Alexa647, Alexa750; 1:2.000 in PBS; Invitrogen, Grand Island, NY, USA and goat secondary antibody conjugated with Cy5, 1:1000 in PBS, Biozol) were incubated for 2 h at RT in the dark. Detection was executed with the fully automated epifluorescence slide scanner microscope AxioScan.Z1 (Zeiss, Oberkochen, Germany) equipped with a Colibri 7 LED system, a Plan-Apochromat 20x/0.8 objective and appropriate beam splitters for DAPI (405 nm), Alexa488 (495 nm), Alexa555 (573 nm), Alexa647/Cy5 (652 nm) and Alexa750 (762 nm). Image stacks (5 µm, variance projection) were recorded and analyzed with ZEN blue (Zeiss, Oberkochen, Germany). Confocal magnification images (512×512 pixel, 1.54 µs pixel dwell time) were acquired with the confocal laser-scanning microscope cLSM 880 (Zeiss, Oberkochen, Germany) using a Plan-Apochromat 63x/1.4 Oil DIC M27 objective, argon (488 nm) and helium-neon laser (542 nm) excitation and appropriate beam splitters. The setup and image acquisition were controlled by ZEN black 2.3 software (Zeiss, Oberkochen, Germany).

### Magnetic Cell Separation (MACs)

Magnetic Cell Separation (MACs) of OPCs was performed as previously described (*28*). Briefly, anesthetized animals were perfused using ice-cold Hank’s balanced salt solution without Ca^2+^ and Mg^2+^ (HBSS, Gibco) and the spinal cord was dissected on ice. The sorting was performed on the basis of the manufacturer’s instructions (Miltenyi Biotec) with minor adjustments: First, debris removal (130-107-677) and resuspension in re-expressing medium (NeuroBrew-21, 1:50, Cat. No. 130-093-566 and 200 mM L-glutamine, 1:100, Sigma; in MACs Neuro Medium, Cat. No. 130-093-570) at 37 °C for 30 min were performed. Subsequently, myelin removal (130-096-731) for 15 min at 4 °C and incubation in Fc-receptor blocker for 10 min at 4 °C were performed. Finally, cells were incubated at 4 °C with 10 µl microbead mixture with antibodies against CD140 (130-101-502), NG2 (130-097-170) and O4 (130-096-670) in a 1:1:1 ratio. MACs-sorted OPCs and flow samples were lysed in RIPA buffer (Thermo Fisher Scientific) and stored at −80 °C until further analysis.

### Real-time PCR

For quantitative real-time PCR (qRT-PCR), mRNA was extracted using the NucleoSpin RNA Plus XS kit (Macherey-Nagel) and cDNA was produced using the Omniscript kit (QIAGEN). qRT-PCR was performed using the EvaGreen kit (Axon) with a CFX96 Real Time System (BioRad). The primer sequences used are listed in Supplementary Table 1.

### Software

For 2P-LSM acquisition, the open-source MATLAB-based software application ScanImage® (Vidrio Technologies, Ashburn, VA, USA) (*54*) was used. The custom-made MATLAB-based software MSparkles, GraphPad Prism 9 and Microsoft Office Excel 2019 were used for data analysis. Immunohistochemical data were collected, visualized and modified using the ZEN imaging software (Zeiss, Oberkochen, Germany) and the ImageJ collection Fiji. For figure layout, the Adobe Creative Suite 2023 was used (Adobe InDesign®, Adobe Illustrator®, Adobe Photoshop®). Administration of the animals was performed with PyRAT, a Python based Relational Animal Tracking software (Scionics Computer Innovation; Dresden, Germany).

### Statistics

Unless otherwise stated, data were analyzed using a Shapiro-Wilk normality test and accordingly represented as mean ± SEM or median with interquartile range of single mice. Single datasets belonging to the same animal were similarly analyzed and represented as mean or median. Data were compared using appropriate parametric or non-parametric tests in line with the result of the Shapiro-Wilk normality test. Test details are given in the corresponding figure legend. For statistical analysis, following *p-values* were used: *, # p<0.05; ** p<0.01; *** p<0.001, **** p<0.0001.

### Ethics statement

All animal experiments were performed at the University of Saarland, Center for Integrative Physiology and Molecular Medicine (CIPMM), in strict accordance with the recommendations to European and German guidelines for the welfare of experimental animals and approved by the “Landesamt für Gesundheit und Verbraucherschutz” of the state of Saarland (animal license number 34/2016, 36/2016, 03/2021 and 08/2021). Number of animals used for each experiment are listed in the corresponding figure legend. In this study, animals of both sexes were used and were equally represented. No sex difference was observed across the study and data were accordingly not segregated into two groups.

## Results

### GABA_B_R deletion in OPCs does not substantially affect OPC physiology in the spinal cord

To investigate the contribution of metabotropic GABA_B_ receptors (GABA_B_R) in oligodendrocyte-precursor cells (OPCs), we induced the time-controlled and cell-specific GABA_B_R deletion in NG2^+^ OPCs coupled to the expression of the genetically-encoded lineage tracer and Ca^2+^ indicator GCaMP3 (GABA_B_R cKO mice). To this aim, 10 week-old mice were injected i.p. for five consecutive days with tamoxifen at a daily dose of 100 mg/kg body weight (Figure 1A), leading to the reduction of the *gabbr1* mRNA levels in sorted OPCs by ∼ 40 % (p = 0.034) (Supplementary Figure 1A-D) and the recombination of more than 80 % of PDGFRα^+^ OPCs two weeks after the first TAM injection (Supplementary Figure 1E-F). Although longer latency between knock-out induction and analysis results in higher efficiency (*48*), earlier tamoxifen administration also resulted in an increased proportion of recombined mature oligodendrocytes as result of the differentiation of recombined OPCs (Supplementary Figure 2), whereas our chosen experimental design reduced the proportion of recombined MyRF^+^ mature oligodendrocyte recombination to ∼ 5-6 % in the gray matter and to ∼ 2-3 % in the white matter (Supplementary Figure 1G).

**Figure 1.**
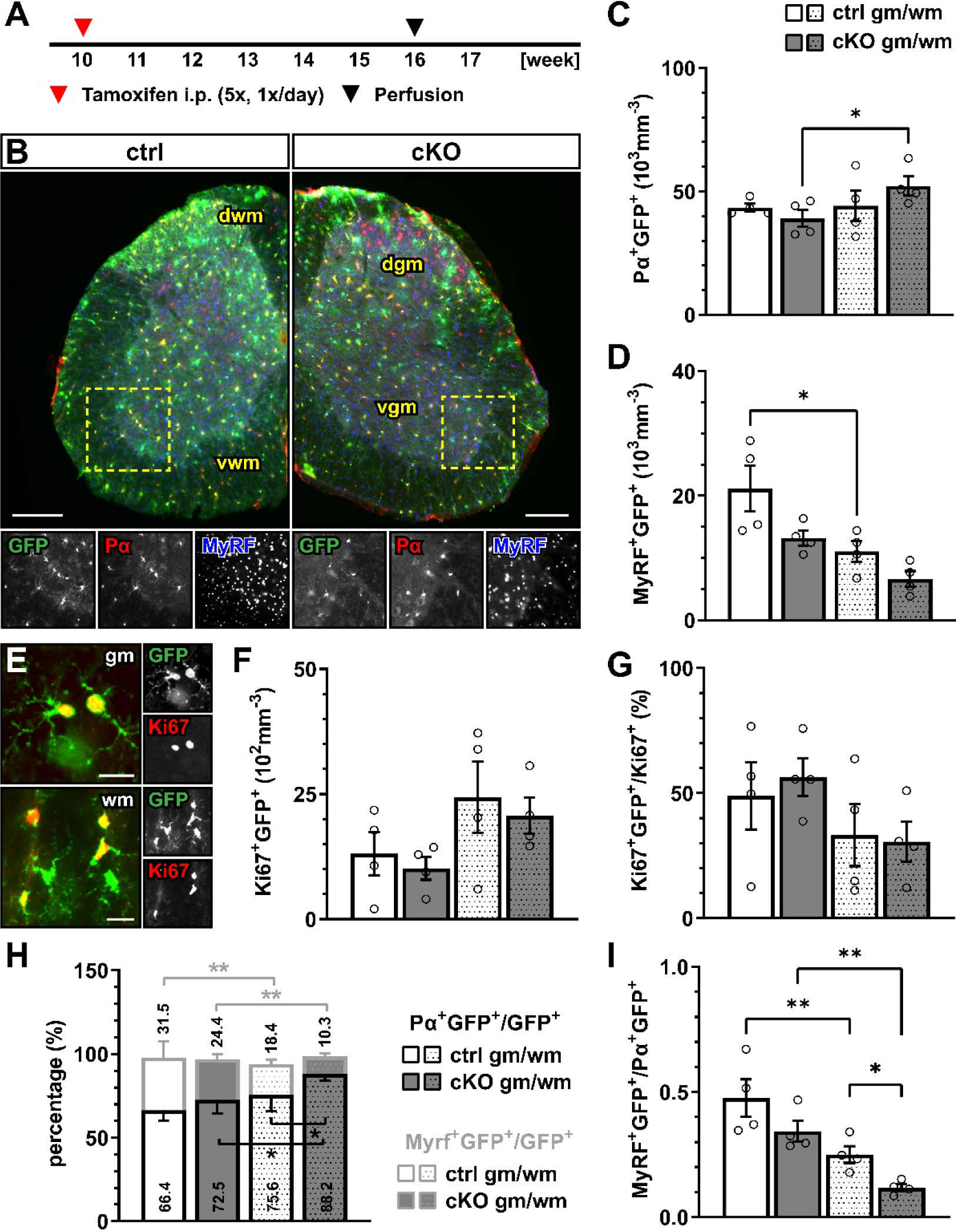
GABA_B_R deletion in OPCs does not affect oligodendrocyte-lineage cell proliferation and differentiation. **(A)** Experimental design for tamoxifen-induced GABA_B_R deletion and GCaMP3 (GFP) expression in NG2^+^ OPCs, perfusion and immunohistochemical analysis (IHC). (**B**) IHC of spinal cord tissue from control (ctrl) and conditional knock-out (cKO) mice stained for GFP (green), PDGFRα (Pα, red) and MyRF (blue). dwm, dorsal white matter; dgm, dorsal gray matter; vgm, ventral gray matter; vwm, ventral white matter. Scale bar, 200 µm. Recombined OPC (Pα^+^GFP^+^, **C**) and mature oligodendrocyte (MyRF^+^GFP^+^, **D**) cell density. (**E**) IHC analysis of proliferating cells (Ki67^+^, red). Scale bar, 20 µm). Recombined proliferating cell density (**F**) and proportion of recombined cells (Ki67^+^GFP^+^) in the total pool of proliferating cells (Ki67^+^) (**G**). (**H**) Proportion of recombined OPCs and mature oligodendrocytes in the total pool of recombined cells. (**I**) Ratio between recombined oligodendrocytes and recombined OPCs. Data are represented as mean ± SEM and derive from N=3-4 mice (n=12 FOVs). Data were analyzed using a Two-way ANOVA with multiple comparisons test.

In the gray and white matter of lumbar spinal cord tissue, the genetic removal of GABA_B_R from OPCs did not affect the density of recombined PDGFRα^+^ OPCs (Figure 1A-C) or the density of recombined MyRF^+^ mature oligodendrocytes (Figure 1D) the between control (ctrl) and GABA_B_ conditional knock-out mice (cKO). Similarly, non-recombined cells did not differ between the two groups (Supplementary Figure 3). In line with these results, no overall alteration of the proliferation rate was detected using the proliferation nuclear marker Ki67 (Figure 1E-F). Finally, to evaluate the differentiation of recombined OPCs, we quantified the percentage of recombined OPCs (PDGFRα^+^GFP^+^) and recombined oligodendrocytes (MyRF^+^GFP^+^) on the total number of recombined cells (GFP^+^). Specifically in the white matter, the relative number of PDGFRα^+^GFP^+^ was increased in cKO mice (ctrl, 75.62 ± 4.95 %; cKO, 88.20 ± 2.08 %; p = 0.011), suggesting a reduced OPC differentiation in cKO mice (Figure 1H). On the other hand, the relative number of MyRF^+^GFP^+^ did not differ between ctrl and cKO mice. The increased PDGFRα^+^GFP^+^ relative number in the white matter of cKO mice was reflected by a reduced MyRF^+^GFP^+^/PDGFRα^+^GFP^+^ ratio representing the number of mature oligodendrocytes produced by a single recombined OPC (Figure 1I).

Next, we investigated whether OPC-GABA_B_R cKO mice display any myelin alteration in the white matter of the spinal cord (Figure 2A-B). No difference was detected in terms of the mean fluorescence intensity of the myelin basic protein (MBP) (Figure 2C) as well as in terms of the myelinated area (Figure 2D). To exclude that these results mask underlying subtle differences in axonal myelination we performed higher-magnification confocal acquisition of MBP, coupled with the immunohistochemical detection of neurofilament positive neuronal axons (Figure 2E). The results showed no difference in axon number (Figure 2F) as well as average axon diameter (Figure 2G). Accordingly, we estimated the mean axonal g-ratio (as the ratio of the inner axonal diameter to the total outer diameter) and its distribution and found no difference between ctrl and cKO mice (Figure 2H).

**Figure 2.**
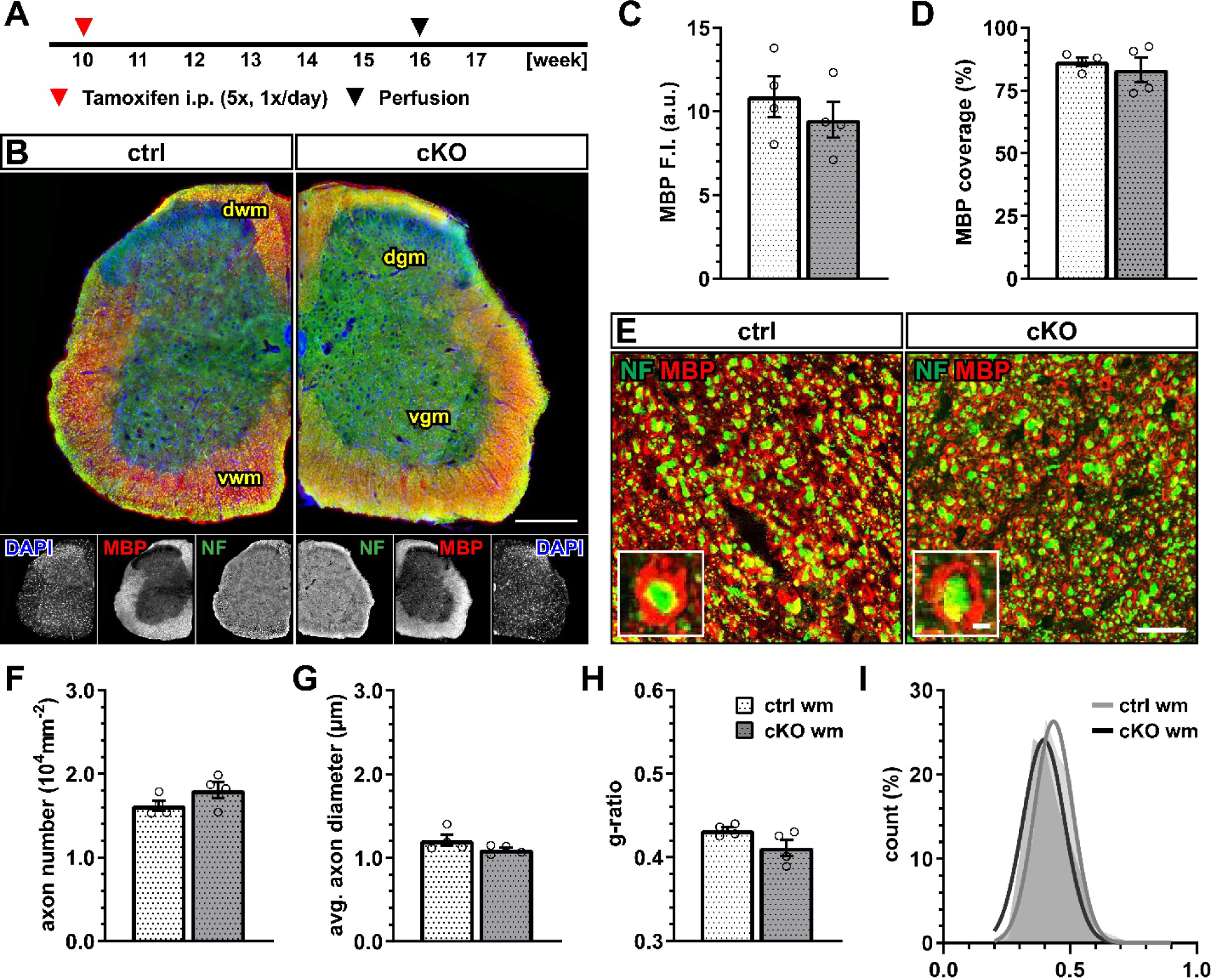
No changes of spinal white matter myelination after GABA_B_R loss. **(A)** Experimental design for tamoxifen-induced GABA_B_R deletion and GCaMP3 (GFP) expression in NG2^+^ OPCs, perfusion and IHC. (**B**) IHC of spinal cord tissue from ctrl and cKO mice stained for DAPI (blue), MBP (red) and neurofilament (green). Scale bar, 200 µm. (**C**) Mean fluorescence intensity (F.I.) of white matter MBP signal and (**D**) MBP coverage expressed as the percentage of the MBP^+^ area in selected FOVs. (**E**) Confocal analysis of white matter spinal cord tissue stained for MBP (red) and neurofilament (green). Scale bar, 10 µm. (**F**) Axonal density and (**G**) average axonal diameter. (**H**) Estimated average axonal g-ratio and (**I**) histogram of the g-ratio distribution (as filled area) as well as non-linear Gaussian fit (as solid line). Data are represented as mean ± SEM and derive from N=4 mice (n=16 FOVs). In **H** and **I**, data derive from n=160 axons per group. Data were analyzed using an unpaired *t* test. In **I**, non-linear fitting was performed using a Least-Squares fitting with no weighting method and compared using the extra-sum-of-squares F test.

These data therefore suggest that the genetic removal of OPC-GABA_B_Rs is not associated with obvious alterations in the physiological level of cell proliferation and differentiation of the oligodendrocyte-lineage as well as myelination within the spinal cord.

### OPCs display increased differentiation upon cuprizone treatment after OPC-GABA_B_R loss

The cuprizone model is a copper-chelating toxic model used to induce demyelination in the CNS mimicking several aspects of human multiple sclerosis (MS), including demyelination, oligodendrocyte death, astrogliosis, microgliosis, OPC proliferation and maturation (*58, 59*). The cuprizone-induced response in the spinal cord has been reported to be reduced compared to the other regions of the forebrain (*60–62*). In virtue of the previous reports on the applicability of the model in the spinal cord, we modified the previously used 0.2 % w/w cuprizone powder feeding model and fed 12-week old mice with a dose of 0.3 % w/w cuprizone for one week and with a dose of 0.2 % w/w for the two following weeks (Figure 3A).

**Figure 3.**
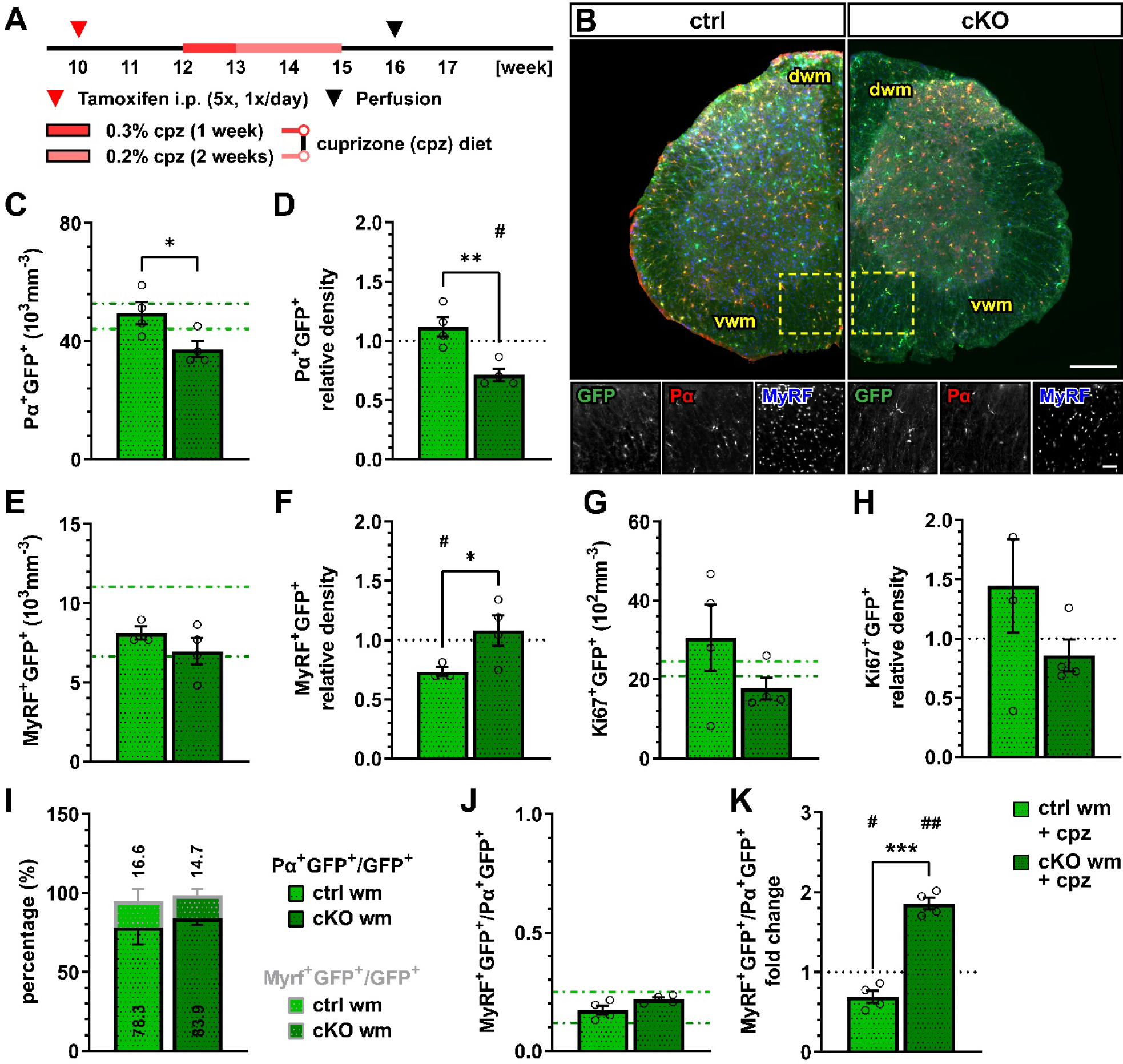
OPC-GABA_B_R loss leads to reduced OPC density and unaltered mature oligodendrocyte density after cuprizone treatment. **(A)** Experimental design for tamoxifen-induced GABA_B_R deletion and GCaMP3 (GFP) expression in NG2^+^ OPCs, cuprizone (cpz) treatment, perfusion and IHC analysis. (**B**) IHC from cuprizone-treated ctrl and cKO mice stained for GFP (green), PDGFRα (Pα, red) and MyRF (blue). Scale bar, 200 µm (overview), 100 µm (zoom-in). Recombined OPC (Pα^+^GFP^+^, **C**) cell density and fold-change (**D**) compared to untreated ctrl and cKO mice. Recombined mature oligodendrocyte (MyRF^+^GFP^+^, **E**) cell density and fold-change (**F**). Recombined proliferating (Ki67^+^GFP^+^, **G**) cell density and fold-change (**H**). (**I**) Proportion of recombined OPCs and mature oligodendrocytes in the total pool of recombined cells. (**J**) Ratio between recombined oligodendrocytes and recombined OPCs and (**K**) fold-change compared to untreated ctrl and cKO mice. Data are represented as mean ± SEM and derive from N=3-4 mice (n=12 FOVs). In **C**, **E** and **G**, data from untreated groups are overlapped as dotted lines (ctrl, light green; cKO, dark green). Data were analyzed using an unpaired *t* test (**C-I**, **K**) or a two-way ANOVA with multiple comparisons test (**J**). For each group in **D**, **F** and **H**, data were tested using a one sample *t* test to compare their mean with the hypothetical value μ=1.

One week after withdrawal of the cuprizone diet, we performed immunohistochemical analysis of the oligodendrocyte-lineage (Figure 3B). Recombined OPC density in the white matter of cuprizone-treated OPC-GABA_B_R cKO mice (37.24 ± 2.73 x10^3^ mm^-3^) was reduced compared to cuprizone-treated controls (49.49 ± 3.71 x10^3^ mm^-3^, p=0.031) (Figure 3C). Also, compared to untreated control mice, cuprizone-treated controls showed no change in the density of recombined OPCs, whereas cKO mice displayed ∼ 30 % decrease in OPC cell density (0.71 ± 0.05, p=0.012) (Figure 3D). Interestingly, the cell density of non-recombined OPCs was increased in cuprizone-treated OPC-GABA_B_R cKO mice compared to cuprizone-treated control mice (Supplementary Figure 4A-C) as well as to untreated cKO mice (Supplementary Figure 4D-E), suggesting that non-recombined OPCs may proliferate to compensate for the reduced density of GABA_B_R-lacking OPCs.

No difference was observed in terms of recombined mature oligodendrocyte cell density between cuprizone-treated control (8.12 ± 0.43 x10^3^ mm^-3^) and cKO mice (6.97 ± 0.82, p=0.628) (Figure 3E). To note though, only control mice showed a reduction in MyRF^+^GFP^+^ cell density upon cuprizone treatment (0.73 ± 0.04, p=0.021), whereas OPC-GABA_B_R cKO mice did not show any difference compared to untreated animals (1.05 ± 0.12, p=0.723) (Figure 3F). These data suggest that the removal of GABA_B_Rs from the oligodendrocyte-lineage cells may be associated with enhanced OPC differentiation upon cuprizone treatment resulting in increased mature oligodendrocyte density. Interestingly, the density of non-recombined mature oligodendrocytes in cKO mice was reduced by the cuprizone treatment similarly to control mice (Supplementary Figure 4D-E), indicating that the protective effect of the genetic removal of OPC-GABA_B_Rs is restricted to recombined cells and their progeny.

With respect to cell proliferation, the density of recombined Ki67^+^ cells, indicative of proliferating recombined OPCs, did not differ either between cuprizone-treated control and cKO mice (Figure 3G) or between treated and untreated groups (Figure 3H). To note, no difference was detected in the relative number of PDGFRα^+^GFP^+^ or MyRF^+^GFP^+^ cells (Figure 3I) and the MyRF^+^GFP^+^/PDGFRα^+^GFP^+^ ratio between controls and cKO mice (Figure 3J). On the other hand, the MyRF^+^GFP^+^/PDGFRα^+^GFP^+^ ratio was almost doubled in cKO mice compared to untreated mice (1.86 ± 0.07, p=0.001), whereas it was reduced in control mice (0.69 ± 0.08, p=0.029) (Figure 3K), in line with enhanced OPC differentiation after loss of the OPC-GABA_B_R.

Astro- and microgliosis in response to cuprizone treatment was evaluated by means of immunohistochemical analysis of the glial fibrillary acidic protein (GFAP) and the ionized calcium-binding adapter molecule 1 (Iba1), respectively (Supplementary Figure 5A-B). In line with the reduced effect of cuprizone on spinal cord tissue published before, no clear white matter astroglial activation or difference between control and cKO mice was detected upon cuprizone treatment (Supplementary Figure 5C-D). The same was true for the number of Iba1^+^ cells (Supplementary Figure 5E), which did not increase upon cuprizone treatment nor was different between the groups (Supplementary Figure 5F), further highlighting the subtle effect of CPZ in the spinal cord.

Given the enhanced number of oligodendrocytes and the increased OPC differentiation upon cuprizone treatment, these results suggest that the deletion of GABA_B_Rs from OPCs protects against demyelination in the spinal cord.

### Cuprizone-induced myelin loss in the spinal cord is ameliorated in OPC-GABA_B_Rs cKO mice

Next, we evaluated the effect of the cuprizone on myelin in control and OPC-GABA_B_R cKO mice (Figure 4A-B). Although the mean MBP fluorescence intensity was only slightly enhanced in cKO compared to control mice (ctrl, 7.80 ± 0.65; cKO, 9.10 ± 0.94; p=0.371) (Figure 4C), MBP was reduced compared to untreated conditions specifically in control mice (0.72 ± 0.06, p=0.018), but unaltered in cKO mice (0.96 ± 0.10, p=0.703) (Figure 4D). Similar considerations hold true for the MBP coverage (ctrl, 65.19 ± 4.99; cKO, 73.56 ± 7.03; p=0.260) (Figure 4E), which was reduced in control mice (0.76 ± 0.07, p=0.036) and left unaltered in cKO mice (0.88 ± 0.05, p=0.107) (Figure 4F).

**Figure 4.**
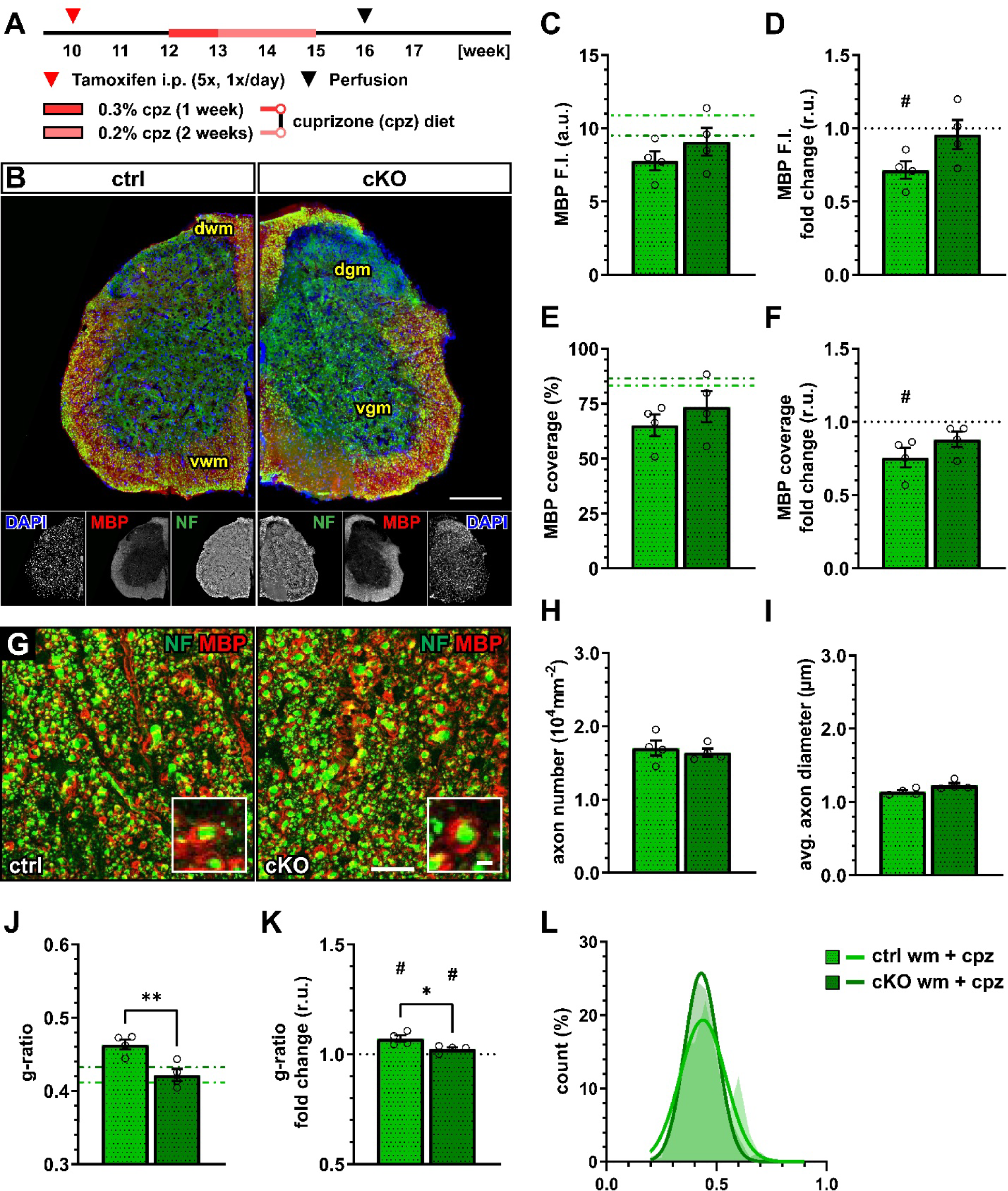
OPC-GABA_B_R cKO protects against cuprizone-induced myelin loss in the spinal cord. **(A)** Experimental design for tamoxifen-induced GABA_B_R deletion and GCaMP3 (GFP) expression in NG2^+^ OPCs, cuprizone (cpz) treatment, perfusion and IHC. (**B**) IHC of spinal cord tissue from ctrl and cKO mice stained for DAPI (blue), MBP (red) and neurofilament (green). Scale bar, 200 µm. (**C**) Mean fluorescence intensity (F.I.) of white matter MBP signal and (**D**) fold-change compared to untreated ctrl and cKO mice. (**E**) MBP coverage expressed as the percentage of the MBP^+^ area in selected FOVs and (**F**) fold-change compared to untreated ctrl and cKO mice. (**G**) Confocal analysis of white matter spinal cord tissue stained for MBP (red) and neurofilament (green). Scale bar, 10 µm. (**H**) Axonal density and (**I**) average axonal diameter. (**J**) Estimated average axonal g-ratio, (**I**) fold-change and (**L**) histogram of the g-ratio distribution (as filled area) as well as non-linear Gaussian fit (as solid line). Data are represented as mean ± SEM and derive from N=4 mice (n=16 FOVs). In **J-K**, data derive from n=160 axons per group. In **C**, **E** and **J** data from untreated groups are overlapped as dotted lines (ctrl, light green; cKO, dark green). Data were analyzed using an unpaired *t* test. For each group in **D**, **F** and **K**, data were tested using a one sample *t* test to compare their mean with the hypothetical value μ=1. In **L**, non-linear fitting was performed using a Least-Squares fitting with no weighting method and compared using the extra-sum-of-squares F test.

Confocal analysis of myelinated axons (Figure 4G) showed no difference in terms of axon density (Figure 4H) as well as average axonal diameter (Figure 4I), indicating no difference in axonal survival or structure between control and cKO mice and their response to the toxicity of cuprizone. On the other hand, the estimated mean axonal g-ratio in cKO mice (0.42 ± 0.01) was lower than under control conditions (0.46 ± 0.01; p=0.002) indicating higher myelination in the cKO mice after cuprizone treatment (Figure 4J) in line with the higher MBP fluorescence. Compared to untreated conditions, both groups displayed an increased g-ratio (ctrl: 1.07 ± 0.01, p=0.016; cKO: 1.03 ± 0.01, p=0.050) but the increase in cKO mice was lower than control mice (p=0.033) (Figure 4K-L). These data are therefore in line with a protective role of OPC GABA_B_R loss to the pathological alteration of myelin quality associated to the cuprizone treatment.

### Also acute myelin loss upon lysolecithin incubation is reduced in OPC-GABA_B_R cKO mice

To confirm the protective effect of OPC-GABA_B_R loss on demyelinating insults, we took advantage of an acute model of demyelination, namely the incubation with the gliotoxic membrane-dissolving compound lysolecithin (or lysophosphatidylcholine, LPC) (*52, 63*). Focal timely-controlled application of LPC on exposed sections of the lumbar spinal cord was used to evaluate the effect of acute demyelinating insults on the spinal dorsal white matter in control and cKO mice (Figure 5A). Contrary to the cuprizone model, LPC directly acts on immune cells, being a direct activator for monocytes, macrophages and lymphocytes (*64–66*). Therefore, inflammation is a predominant component of the model next to the direct detergent effect of LPC on oligodendrocytes. In line with this, we detected a strong glial response upon LPC incubation on the superficial layers of the dorsal white matter tracts in control and cKO mice (Supplementary Figure 6A-B), both in terms of astroglial GFAP expression (Supplementary Figure 6C-D) and Iba1^+^ microglia activation (Supplementary Figure 6E-F).

**Figure 5.**
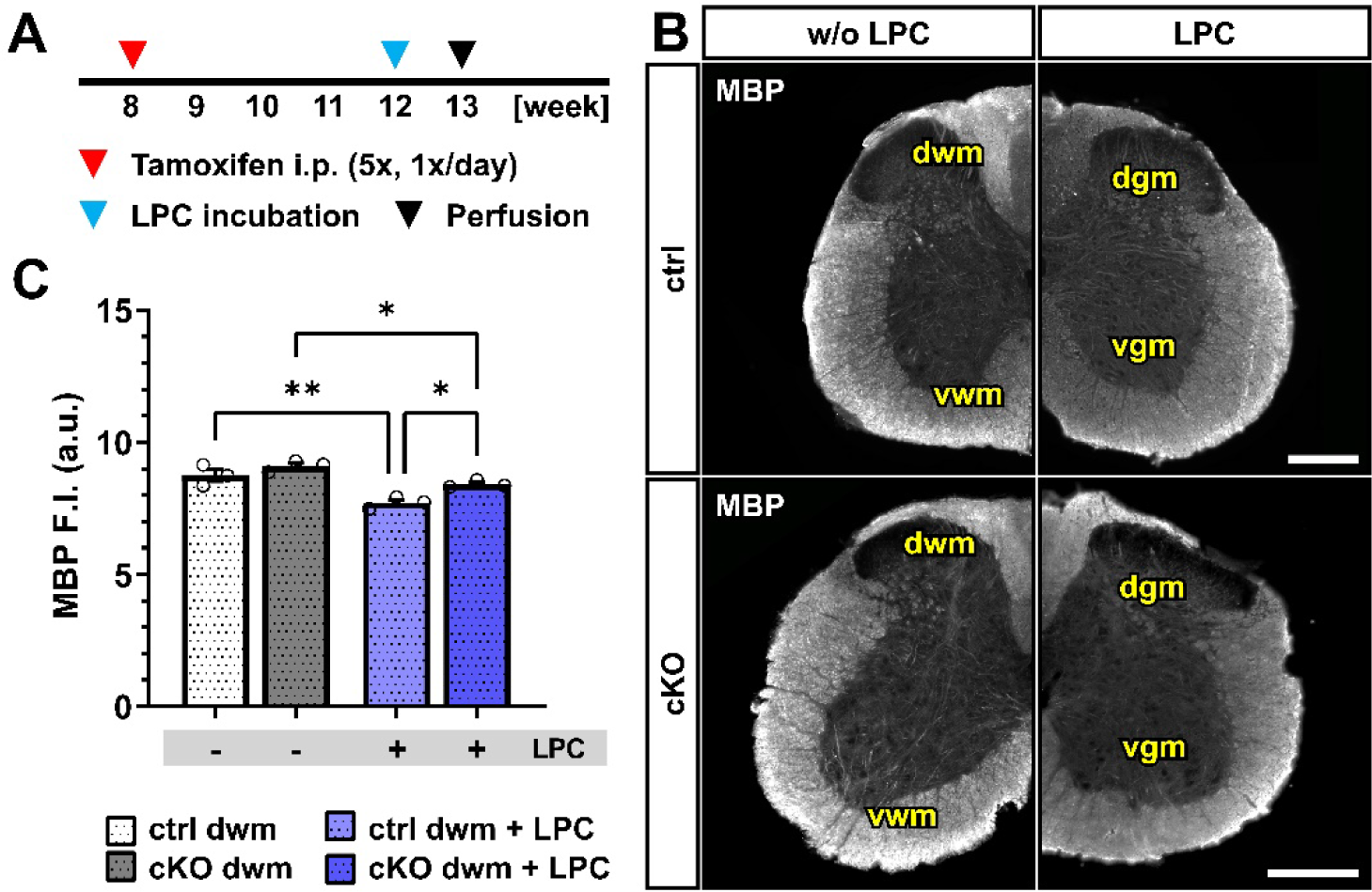
Acute myelin loss upon lysolecithin incubation reduced in OPC-GABA_B_R cKO mice. **(A)** Experimental design for tamoxifen-induced GABA_B_R deletion and GCaMP3 (GFP) expression in NG2^+^ OPCs, lysolecithin (LPC) incubation, perfusion and IHC. (**B**) IHC of spinal cord tissue from ctrl and cKO mice stained for MBP. Scale bar, 200 µm. (**C**) Mean fluorescence intensity (F.I.) of white matter MBP signal. Data are represented as mean ± SEM and derive from N=3 mice (n=12 FOVs). In **J-K**, data derive from n=160 axons per group. Data were analyzed using an unpaired *t* test. Data were analyzed using a Two-way ANOVA with multiple comparisons test.

Upon LPC incubation, control mice showed a clear reduction in the mean MBP fluorescence intensity restricted to the dorsal white matter (w/o LPC, 8.75 ± 0.23; LPC, 7.71 ± 0.13; p=0.001). OPC-GABA_B_R cKO mice also were associated with a MBP reduction (w/o LPC, 9.11 ± 0.12; LPC, 8.4 ± 0.08; p=0.011) (Figure 5B-C) but to a lesser extent compared to LPC-treated control mice (p=0.011). As it was the case for the cuprizone model, the data support a protective role of the cKO of OPC-GABA_B_Rs to demyelinating insults.

### Spinal cord OPCs display altered Ca^2+^ dynamics in OPC-GABA_B_R cKO mice

Accumulating evidence revealed that Ca^2+^ signaling is a key regulator of OPC proliferation and differentiation (*39, 40*) as well as myelination (*35–37, 67*). In particular, it has been recently suggested that OPC Ca^2+^ activity reduces along their differentiation to mature oligodendrocytes (*40*). In virtue of these considerations, and given the putative role of GABA_B_Rs on regulation of intracellular Ca^2+^ concentration and signaling (*22*), we used in vivo two-photon laser scanning microscopy (2P-LSM) to record spontaneous OPC Ca^2+^ signaling of the dorsal spinal cord white matter tracts to evaluate if the differences in oligodendrocyte-lineage cells and the differential response to demyelination in control and cKO mice could be due to underlying differences in OPC Ca^2+^ (Figure 6A-B). OPC-GABA_B_R cKO mice displayed a drastic increase in the number of recorded Ca2+ oscillations with three times more signals than controls (ctrl, 17.78 ± 3.81 10^-2^µm^-2^min^-1^; cKO, 57.80 ± 6.12 10^-2^µm^-2^min^-1^; p<0.001) (Figure 6C). The increased signal number resulted from an increased spatial signal density, as the regions-of-activity (ROAs) detected in the field-of-view showed a 1.5-fold increase in density (ctrl, 24.16 ± 2.74 10^-3^µm^-2^min^-1^; cKO, 34.21 ± 3.76 10^-3^µm^-2^min^-1^; p=0.007) (Figure 6D), as well as from an increased temporal density, as the per-ROA signal frequency also displayed a 1.6-fold increase (ctrl, 3.86 ± 0.68 min^-1^; cKO, 6.29 ± 0.41 min^-1^; p=0.002) (Figure 6E). On the other hand, the ROA area of the automatically detected Ca^2+^ signals was reduced by ∼ 57 % in cKO mice (ctrl, 82.80 ± 14.92 µm^2^; cKO, 35.26 ± 6.37 µm^2^; p=0.003) (Figure 6F), indicating that the increased Ca^2+^ activity in cKO mice is paralleled by the reduction of its spatial extension. Finally, we did not detect any difference in the signal amplitude between ctrl and cKO mice (Figure 6G) but a slight reduced signal duration in the cKO group (ctrl, 4.98 ± 1.00 s; cKO, 3.17 ± 0.06 s; p=0.044) (Figure 6H). This suggests that the genetic removal of GABA_B_Rs in OPCs impacts the spatial and temporal distribution of OPC Ca^2+^ signaling, resulting in increased Ca^2+^ oscillations with reduced spatial extension and duration.

**Figure 6.**
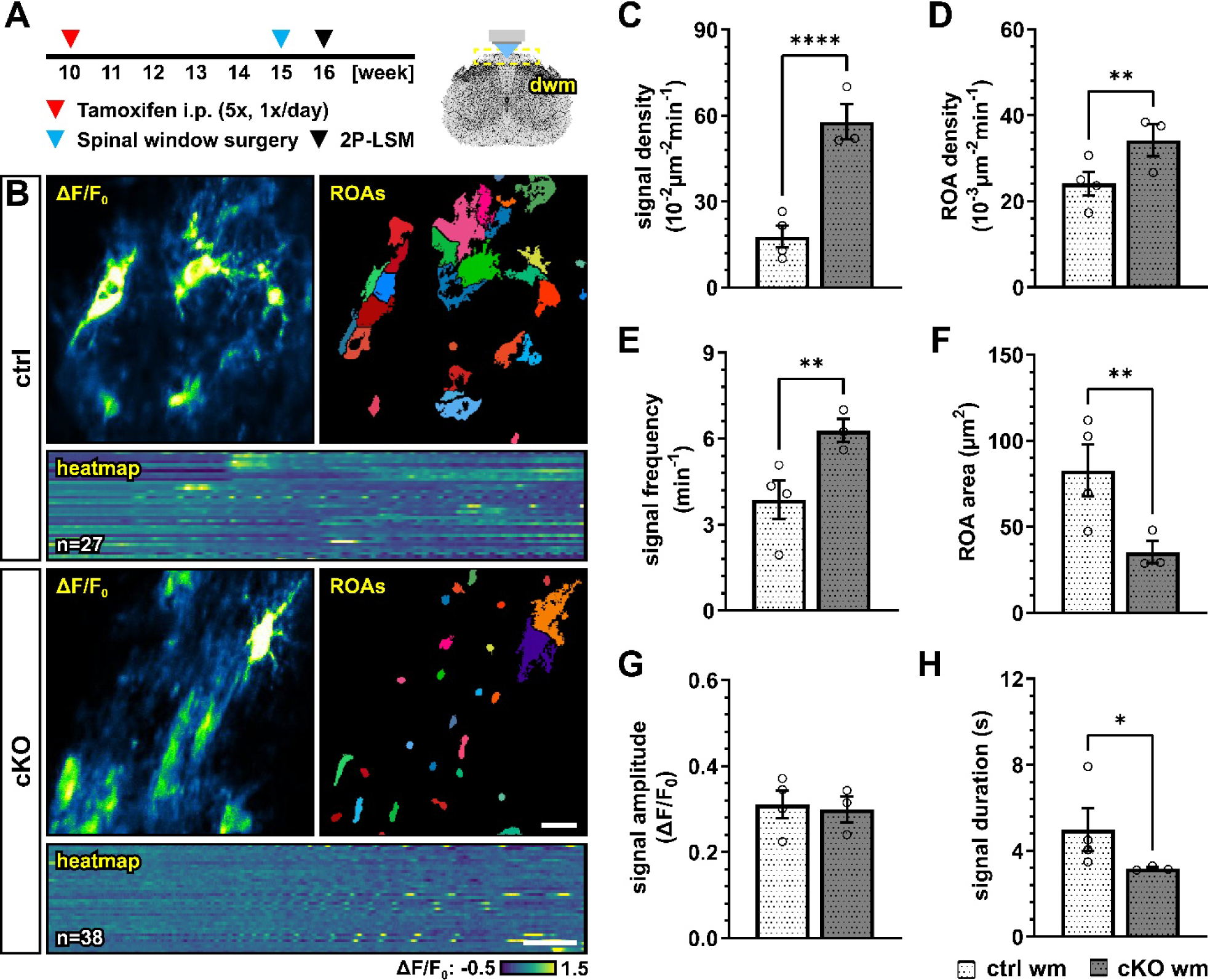
GABA_B_R cKO mice display aberrant Ca^2+^ dynamics in OPCs with increased signal frequency and density but reduced spatial and temporal profiles. **(A)** Experimental design for tamoxifen-induced GABA_B_R deletion and GCaMP3 expression in NG2^+^ OPCs, spinal cord window implantation surgery and *in vivo* two-photon laser-scanning microscopy (2P-LSM). (**B**) Representative maximum-intensity projections of the GCaMP3 signal (ΔF/F_0_) for representative FOVs over the entire recording time (up to 5 min), associated maps of selected identified regions-of-activity (ROAs) with heatmap representation of the GCaMP3 signal profile over time. Scale bar, 20 µm; 30 s. (**C**) Signal density and (**D**) ROA density expressed as the number of signals/ROAs detected over the entire recording time. (**E**) Per-ROA signal frequency. (**F**) ROA area. (**G**) Signal amplitude in correspondence of the signal maximum. (**H**) Signal duration expressed as Full Width at Half Maximum (FWHM). Data are represented as mean ± SEM and derive from N=3-4 mice (n=12 FOVs). Data were analyzed using an unpaired *t* test.

### OPC-GABA_B_R loss counters cuprizone-induced alterations in OPC Ca^2+^ signaling

We next investigated how the cuprizone treatment alters OPC Ca^2+^ activity in control and OPC-GABA_B_R cKO mice. To this aim, we implanted a spinal cord window on 15-week-old mice immediately at the end of the cuprizone treatment and recorded with 2P-LSM spontaneous Ca^2+^ activity from the white matter tracts one week later (Figure 7A-B). As we observed in the untreated groups, GABA_B_R cKO mice displayed also under pathological conditions increased OPC Ca^2+^ signal density (ctrl, 41.41 ± 9.02 10^-2^µm^-2^min^-1^; cKO, 88.13 ± 7.03 10^-2^µm^-2^min^-1^; p<0.001) (Figure 7C). Compared to the untreated-groups, the OPC Ca^2+^ signal density was increased by at least 1.5-fold, with a larger increase in control mice (ctrl, 2.33 ± 0.51, ns, p=0.079; cKO, 1.53 ± 0.12, p=0.049) (Figure 7D). Similarly, control and GABA_B_R cKO mice were associated with increased ROA density (ctrl, 37.08 ± 3.46 10^-3^µm^-2^min^-1^; cKO, 53.82 ± 3.34 10^-3^µm^-2^min^-1^; p=0.006) (Figure 7E) with a similar 1.5-fold change increment compared to untreated conditions in both groups (Figure 7F). After cuprizone treatment, the signal frequency of the OPC Ca^2+^ activity was similar between control (4.99 ± 0.64 min^-1^) and GABA_B_R cKO mice (6.66 ± 0.50 min^-1^, p=0.082) (Figure 7G), resulting from the specific increase of OPC Ca^2+^ activity in control mice compared to untreated conditions (Figure 7H). On the other hand, also the ROA spatial extension did not show any difference after cuprizone treatment between control and cKO mice (ctrl, 50.36 ± 13.59 µm^2^; cKO, 40.29 ± 9.70 µm^2^; p=0.593) (Figure 7I) resulting from the specific decrease in the ROA area in control mice compared to untreated conditions (ctrl, 0.61 ± 0.16, ns, p=0.097; cKO, 1.14 ± 0.28, p=0.656) (Figure 7J). Finally, signal amplitude or duration after cuprizone treatment were similar between control and cKO mice (Figure 7K-L) as well as compared to untreated mice. Taken together, our data suggest that the cuprizone treatment induces an increase in OPC Ca^2+^ activity and that the cKO of OPC-GABA_B_Rs partially counteracts the pathological shift of OPC Ca^2+^ signaling activity.

**Figure 7.**
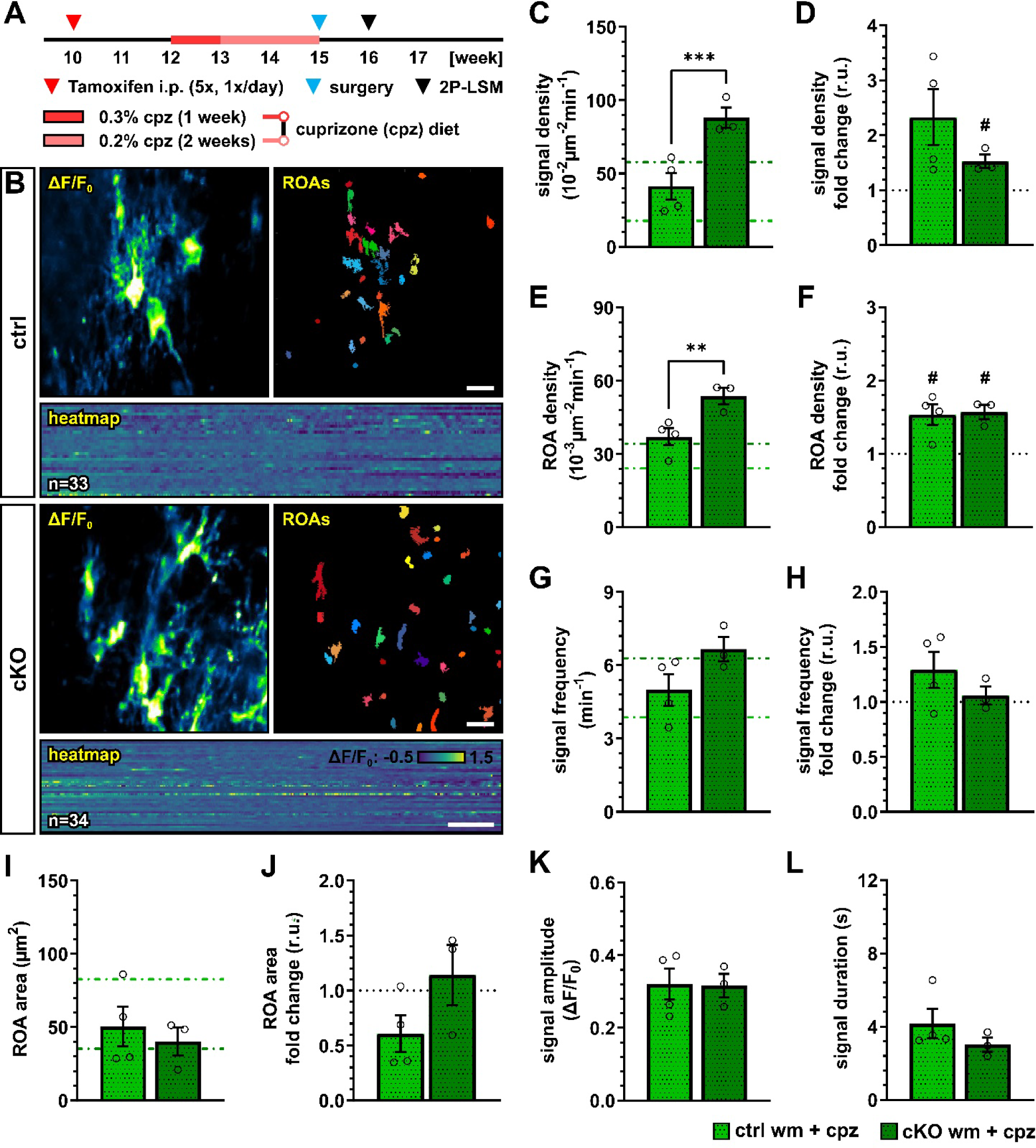
Cuprizone-induced alterations in OPC Ca^2+^ signaling are reduced after OPC-GABA_B_R loss. **(A)** Experimental design for tamoxifen-induced GABA_B_R deletion and GCaMP3 expression in NG2^+^ OPCs, cuprizone treatment, spinal cord window implantation surgery and *in vivo* two-photon laser-scanning microscopy (2P-LSM). (**B**) Representative maximum-intensity projections of the GCaMP3 signal (ΔF/F_0_) for representative FOVs over the entire recording time (up to 5 min), associated maps of selected identified regions-of-activity (ROAs) with associated heatmap representation of the GCaMP3 signal profile over time. Scale bar, 20 µm; 30 s. (**C**) Signal density and (**D**) fold-change compared to untreated ctrl and cKO mice. (**E**) ROA density and (**F**) fold-change. (**G**) Per-ROA signal frequency and (**H**) fold-change. (**I**) ROA area and (**J**) fold-change. (**K**) Signal amplitude and (**L**) signal duration expressed as Full Width at Half Maximum (FWHM). Data are represented as mean ± SEM and derive from N=3-4 mice (n=12 FOVs). In **C**, **E**, **G** and **I** data from untreated groups are overlapped as dotted lines (ctrl, light green; cKO, dark green). Data were analyzed using an unpaired *t* test. For each group in **D**, **F, H** and **J**, data were tested using a one sample *t* test to compare their mean with the hypothetical value μ=1.

## Discussion

In this work, we took advantage of the time- and spatial controlled, cell-type specific GABA_B_R deletion in NG2^+^ OPCs to selectively evaluate their contribution to OPC proliferation, differentiation and maturation in the murine spinal cord. Evidence on the specific role of GABA_B_R signaling in the context of oligodendrocyte-lineage cell physiology is currently limited to *in vitro* studies (*26, 30*), to the brain (*28*) or developmental stages (*32*). Therefore, the transposition of previous results obtained in the field to the spinal cord is challenging, especially in virtue of the high heterogeneity of OPCs across age and CNS location, which has been extensively reported and reviewed (*68–71*). Studies addressing the role of GABA_B_Rs in the spinal cord have been using the GABA_B_R agonist baclofen (*31*). Given the use of baclofen in the therapy of spasticity associated to MS, these results are of key importance. Nevertheless, they still lack the cell-specificity required to identify the specific contribution of OPC-GABA_B_Rs. On the other hand, it is true that both GABA_B1_ and GABA_B2_ subunits are downregulated during OPC differentiation to mature oligodendrocytes (at least *in vitro*) (*30*). Our approach however, selectively targets OPCs with high efficiency, without significantly altering the receptome and physiology of mature oligodendrocytes with the exception of oligodendrocyte newly generated from recombined OPCs (which to the time point of analysis and in the spinal cord never contributed to more than 10 % of existing mature oligodendrocytes) (Supplementary Figure 1E-G).

Our results showed that in the adult spinal cord the conditional OPC specific GABA_B_R knock-out (cKO) does not affect the total number of OPCs and mature oligodendrocytes (in neither the gray nor in the white matter) as it leaves OPC proliferation and differentiation substantially unaltered (Figure 1). It has been recently shown that OPC and OPC-GABA_B_R-signaling are key players in the cortical myelination during development but not in the adult brain (*28*). In line with this, we also did not observe any reduction in spinal myelination in our adult OPC-GABA_B_R cKO mice, neither in terms of MBP coverage nor in terms of myelin thickness (Figure 2). To address the contribution of OPC-GABA_B_Rs in the cellular response to pathological demyelinating insults of the spinal cord, we took advantage of two different models for toxic demyelination, namely the cuprizone and the lysolecithin models. The cuprizone model mimics several aspects of human MS and is associated to a less extensive primary immune response compared to other models including the lysolecithin model (*58, 59*) but its use in the spinal cord has been so far limited due to a reduced response compared to forebrain white matter (*60–62*). Indeed, in line with these observations, we found no specific increase in astrogliosis or microgliosis upon cuprizone treatment (Supplementary Figure 3) as we did not detect any enhanced OPC proliferation in the spinal cord. Nevertheless, the cuprizone treatment effectively reduced the cell density of mature oligodendrocyte in the white matter spinal cord (Figure 3). This result mirrors the higher susceptibility of mature oligodendrocytes (in virtue of their high energy demand) to the chopper chelating action of cuprizone (*13, 72*). On the other hand, OPC-GABA_B_R cKO mice showed no reduction in the number of mature oligodendrocytes, in line with a protective role of the genetic removal of OPC GABA_B_Rs, which could be ascribed to enhanced OPC differentiation (Figure 3).

From a clinical point of view, more important than the enhanced oligodendrocyte survival is the subsequent partial rescue of the cuprizone induced demyelination. OPC-GABA_B_R cKO mice showed no reduction in total MBP and no reduction in the MBP coverage, as it was the case for cuprizone-treated control mice. On the other hand, the analysis of the myelin rings wrapping the white matter axons revealed that both control and cKO mice experienced a thinning of the myelin rings but OPC-GABA_B_R cKO mice exhibited thicker rings, hinting again at a higher resistance to cuprizone (Figure 4). Similar results were obtained from the MBP evaluation in the dorsal white matter upon local lysolecithin treatment (Figure 5). Contrary to the cuprizone model, the loss of MBP was associated with a strong astro- and microglial reaction in control and cKO mice (Supplementary Figure 4), suggesting that the lysolecithin model may be preferred to the cuprizone model for studies on demyelination in the spinal cord. Also, this may be due to the different modes of action of cuprizone and lysolecithin, since lysolecithin, but not cuprizone, acts as a direct activator for immune cells (*64–66*). Nevertheless, beyond their differences, in both models the loss of MBP was reduced in OPC GABA_B_R-deficient mice, suggesting that the protective role of OPC GABA_B_R cKO on demyelination may be independent of its specific trigger. Although the results obtained using the lysolecithin model seem in contradiction to what has been shown using the GABA_B_R agonist baclofen using the same model in the spinal cord (*31*), it is worth mentioning that in our study we induced the cell-type specific GABA_B_R removal before the lesion and focused on the demyelination phase of the model, whereas Serrano-Regal et al. activated GABA_B_R globally after the lesion and focused on remyelination. Therefore, our combined results clearly indicate a complex contribution of GABA_B_R signaling in the context of de- and remyelination in the spinal cord.

This is also highlighted by the complex changes in OPC Ca^2+^ signaling under physiological and pathophysiological conditions. The removal of OPC-GABA_B_Rs under physiological conditions resulted in the alteration of OPC Ca^2+^ oscillations, leading to higher spatial incidence and frequency of Ca^2+^ signals but with reduced spatial (area) and temporal extension (duration) (Figure 6). Glial Ca^2+^ signaling is, as it is in other regions of the CNS, finely regulated and cell-specific in the mouse spinal cord (*50*), but we are far from understanding its determinants as well as its role in cell physiology. In the functional spinal cord, OPC Ca^2+^ signals are known to be evoked by neurons and astroglia (*24*) and disturbances in OPC Ca^2+^ (with excessively frequent Ca^2+^ oscillations) has been linked to impaired proliferation during development (*73, 74*). Compared to physiological conditions in cKO mice, OPCs showed higher resilience to the pathological alteration of cellular Ca^2+^ signaling induced by the cuprizone treatment. In fact, control OPCs displayed an increased Ca^2+^ signaling and reduced Ca^2+^ spatial extension (like the cKO under physiological conditions), which was partially counteracted in cKO mice (Figure 7). It has been shown that sustained Ca^2+^ increase resulting from activation of P2X_7_ purinergic receptors in oligodendrocytes leads to oligodendrocyte death and demyelination *in vivo* (*67*) and that low-frequency Ca^2+^ transients with longer duration in oligodendrocytes are associated with myelin sheath shortening (*36, 37*), which are in line with our results obtained in OPCs and support the use of a common Ca^2+^-coded language for the cells of the oligodendrocyte lineage. To date, little is known about the specific contribution of OPC Ca^2+^ in the context of demyelination. Nevertheless, it has been shown in the spinal cord of zebrafish (*39*) that higher Ca^2+^ activity in OPC is associated with enhanced proliferation and reduced differentiation, whereas lower Ca^2+^ activity is associated with enhanced differentiation. Also, it has been recently shown that in the somatosensory cortex of awake-behaving mice the OPC Ca^2+^ activity decreases as they differentiate into oligodendrocytes (*40*). In line with this, we found that OPC Ca^2+^ activity increases upon cuprizone treatment and is associated with a reduced OPC differentiation and that the loss of OPC GABA_B_Rs counteracts cuprizone-induced pathological alteration of OPC Ca^2+^ activity and is associated with enhanced OPC differentiation upon cuprizone-treatment.

Although our results cannot clarify the contribution of OPC Ca^2+^ to the cell fate and myelination, we could show that resilience to pathological alterations of Ca^2+^ in cKO OPCs was associated with higher endurance to demyelinating insults. It is to be noted that the differences between control and cKO OPCs even in physiological conditions may open up different scenarios, including a preexisting OPC alteration which masks cuprizone-induced changes or conversely a preexisting priming of cKO OPCs which could make them more prone to readily react to demyelination insults.

## Conclusion

Summarized, our study evaluated the role of OPC-GABA_B_Rs signaling in oligodendrocyte-lineage cells in the mouse spinal cord. Despite accumulating evidence on the involvement of OPC responses to GABAergic signaling in the CNS homeostasis and response to demyelinating insults, the specific contribution of OPC-GABA_B_Rs *in vivo* remains elusive. Through targeted genetic removal of GABA_B_Rs from OPCs, we showed that their absence does not alter OPC proliferation or differentiation under physiological conditions in the adult spinal cord. However, the conditional knock-out of OPC-GABA_B_Rs leads to distinctive alterations in OPC Ca^2+^ signaling, emphasizing the intricate regulatory role of GABA_B_Rs in the physiology of oligodendrocyte-lineage cells. Furthermore, our examination of the cuprizone and lysolecithin models of toxic demyelination uncovered a protective effect associated with the removal of OPC-GABA_B_Rs. OPC-GABA_B_R cKO mice exhibited enhanced resilience to cuprizone-induced alterations in OPC Ca^2+^ signaling and, more importantly, a partial rescue of mature oligodendrocyte loss and demyelination. These findings not only shed new light on the involvement of GABA_B_Rs in OPC physiology but also hint at potential therapeutic implications in the context of demyelinating disorders. Overall, our study provides a crucial foundation for further unraveling the intricate dynamics of GABA_B_R signaling in spinal cord oligodendrocyte-lineage cells and their response to demyelination.

## Author Contributions

DG organized the data, performed and supervised data analysis, generated the figures and wrote the manuscript. PR performed the experiments, organized the data and performed data analysis. LF performed MACs and qRT-PCR. EB and PR established the LPC model. MS, EDam and EDal generated and analyzed immunohistochemical data. NB contributed to data analysis. XB contributed to experimental design and supported grant application. FK provided structural and financial support for the project. AS provided financial support and supervised the project, reviewed and finalized the manuscript. FK and AS designed the project. All authors approved on the final version of the manuscript.

## Acknowledgments

The authors thank Dr. Wenhui Huang and the members of the Department of Molecular Physiology of the University of Saarland for intellectual input. We are grateful to Babette Fuss (Virgina Commonwealth University, Richmond, Va., USA) for initial cuprizone protocol discussions. We thank Daniel Schauenburg and colleagues for expert mouse maintenance and tamoxifen treatment.

---

**Supplementary Figure 1.**
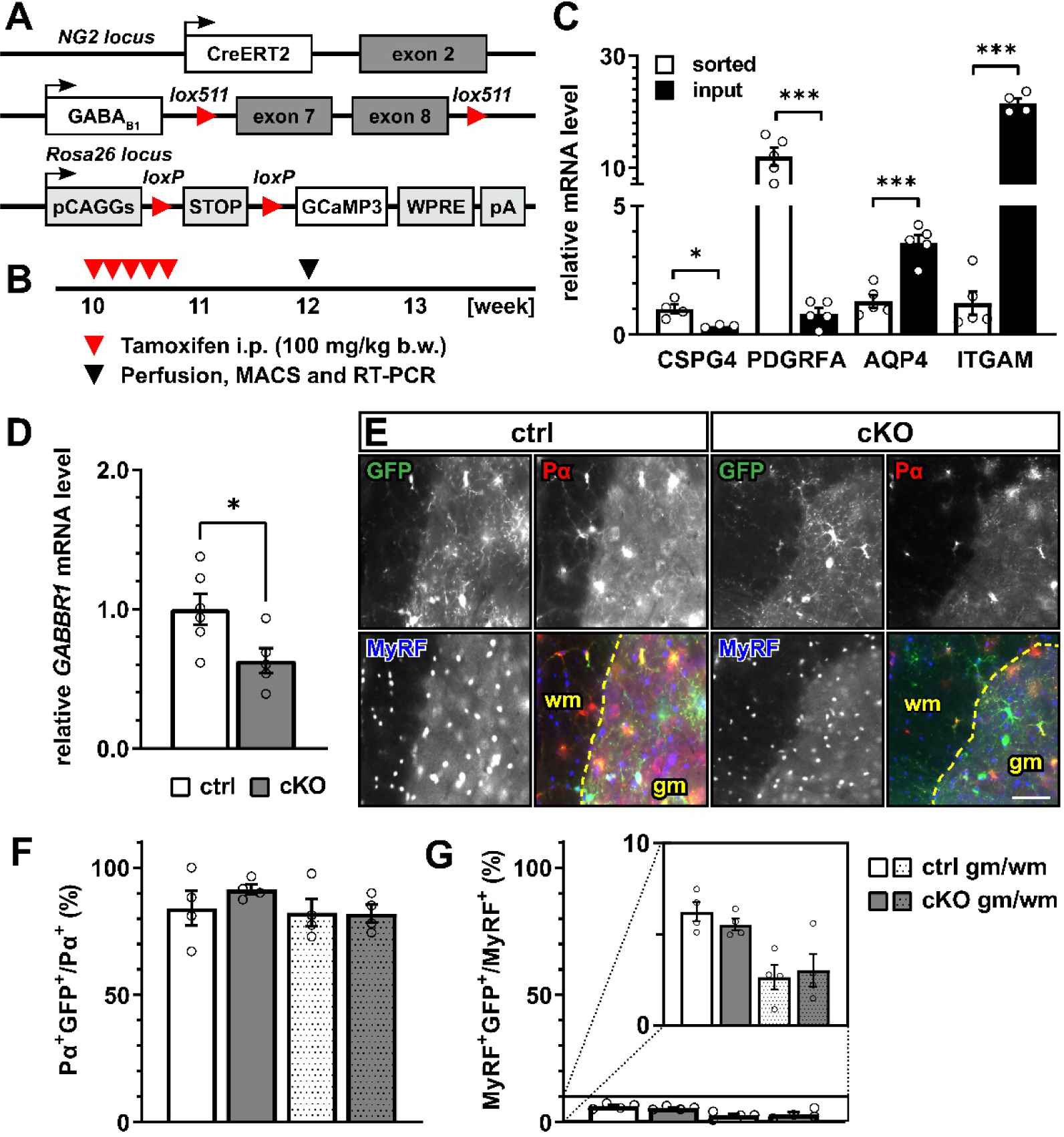
OPC-specific conditional knock-out of GABA_B_Rs. **(A)** Schematic representation of genetic knock-in manipulation required for the tamoxifen-induced GABA_B_R deletion and GCaMP3 (GFP) expression in NG2^+^ OPCs. (**B**) Experimental design for tamoxifen treatment, perfusion, Magnetic Cell Separation Sorting (MACS) and real-time PCR (RT-PCR). (**C**) Cell enrichment analysis of sorted and unsorted (input) cells from spinal homogenate using RT-PCR for OPC-(chondroitin sulfate proteoglycan 4, CSPG4; PDGFRA), astroglial-(aquaporin-4, AQP4) and immune cell-(integrin α M, ITGAM) specific markers. (**D**) RT-PCR of GABBR1 mRNA in spinal homogenates from control (ctrl) and OPC-GABA_B_R conditional knock-out (cKO) mice. (**E**) Immunohistochemistry (IHC) of spinal cord tissue stained for GFP (green), PDGFRα (Pα, red) and MyRF (blue). Scale bar, 20 µm. (**F**) Percentage of recombined OPCs (Pα^+^GFP^+^) on total number of OPCs (Pα^+^). (**G**) Percentage of recombined OPCs (MyRF^+^GFP^+^) on total number of OPCs (MyRF^+^). Data are represented as mean ± SEM and derive from N=3-6 mice for the MACS experiments or N=4 (n=12 FOVs) for IHC. Data were analyzed using a two-way ANOVA with multiple comparisons test (**C**, **F-G**) or an unpaired *t* test (**D**).

**Supplementary Figure 2.**
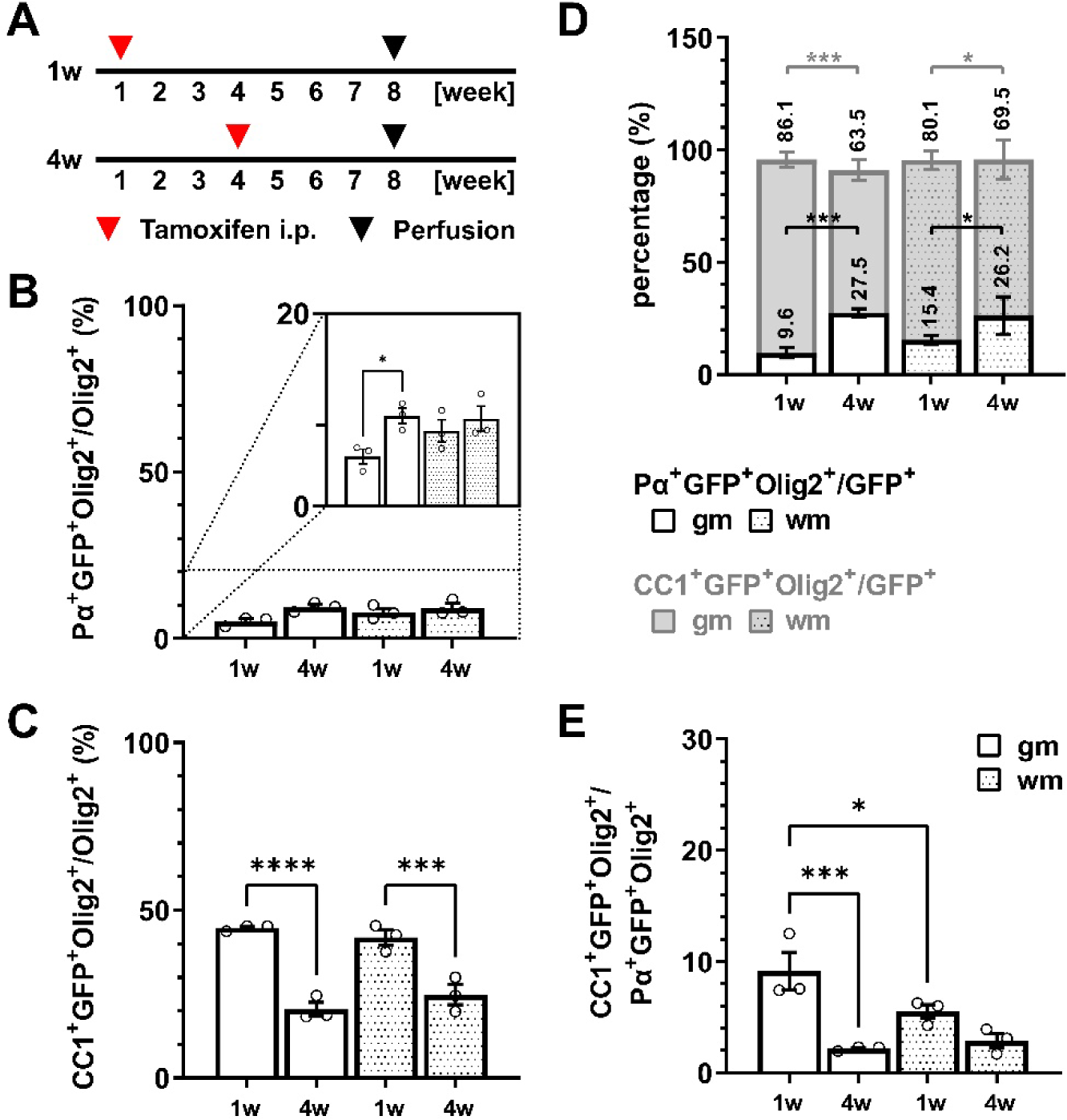
Early tamoxifen administration results in increased number of recombined mature oligodendrocytes. **(A)** Experimental design for tamoxifen-induced GCaMP3 (GFP) expression in NG2^+^ OPCs either at postnatal week one (1w, 2x, 1x/day, i.p.) or postnatal week four (4w, 5x, 1x/day, i.p.). Perfusion and IHC analysis were performed at postnatal week 8. (**B**) Percentage of recombined PDGFRα (Pα)^+^GFP^+^Olig2^+^ OPCs on the total number of Olig2^+^ oligodendrocyte-lineage cells. (**C**) Percentage of recombined CC1^+^GFP^+^Olig2^+^ mature oligodendrocytes on the total number of Olig2^+^ oligodendrocyte-lineage cells. (**D**) Proportion of recombined OPCs and mature oligodendrocytes in the total pool of recombined cells (GFP^+^). (**E**) Ratio between recombined oligodendrocytes and recombined OPCs. Data are represented as mean ± SEM and derive from N=3 mice (n=12 FOVs). Data were analyzed using a Two-way ANOVA with multiple comparisons test.

**Supplementary Figure 3.**
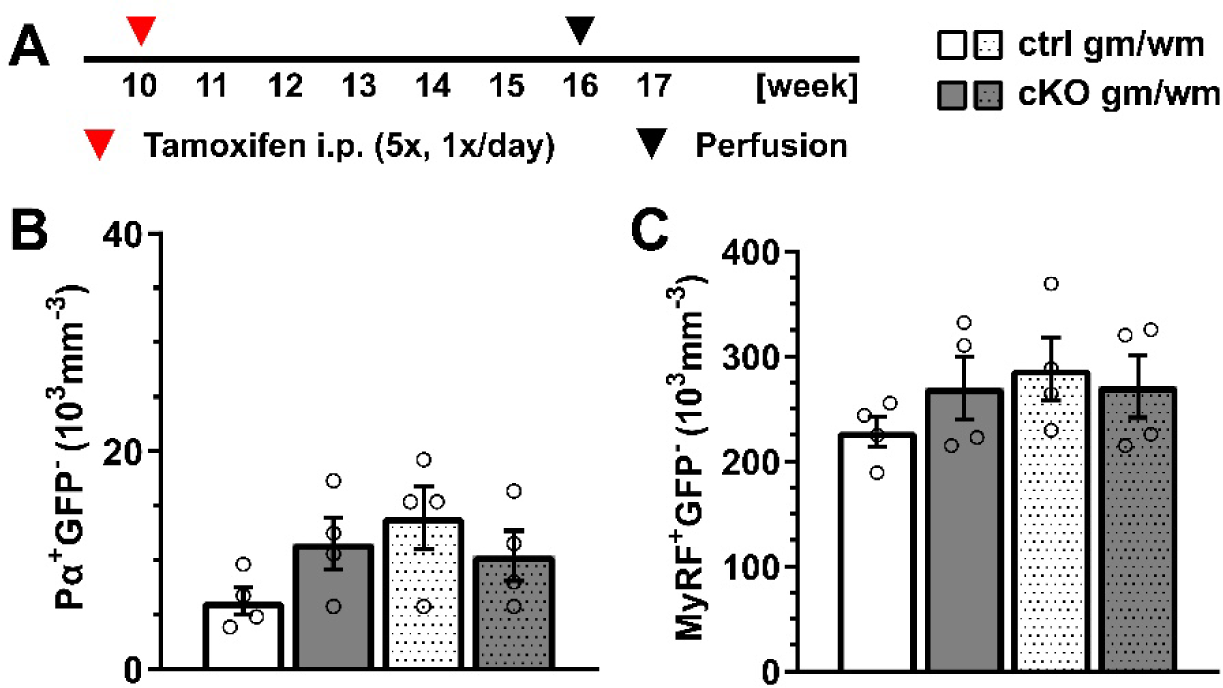
GFP^-^ OPC and mature oligodendrocyte cell density is unaffected in cKO mice. **(A)** Experimental design for tamoxifen-induced GABA_B_R deletion and GCaMP3 (GFP) expression in NG2^+^ OPCs, perfusion and immunohistochemical analysis (IHC). Non-recombined OPC (Pα^+^GFP^-^, **C**) and mature oligodendrocyte (MyRF^+^GFP^-^, **D**) cell density in the gray (gm) and white matter (wm) of the spinal cord. Data are represented as mean ± SEM and derive from N=3-4 mice (n=12 FOVs). Data were analyzed using a Two-way ANOVA with multiple comparisons test.

**Supplementary Figure 4.**
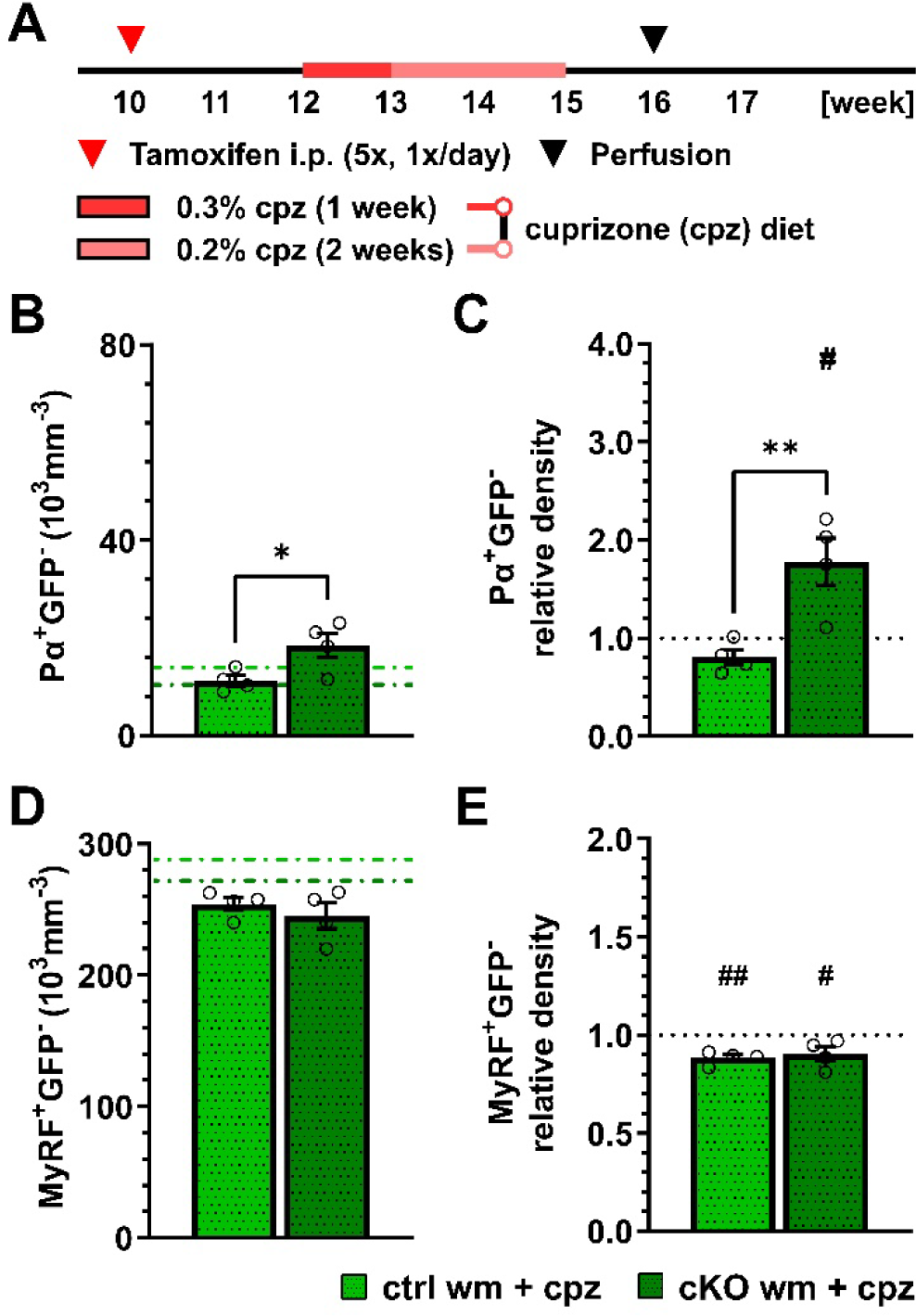
Effect of OPC GABA_B_R loss on cell density of non-recombined oligodendrocyte-lineage cells upon cuprizone treatment. **(A)** Experimental design for tamoxifen-induced GABA_B_R deletion and GCaMP3 (GFP) expression in NG2^+^ OPCs, cuprizone (cpz) treatment, perfusion and IHC analysis. of ctrl and cKO mice. Non-recombined OPC (Pα^+^GFP^-^, **B**) cell density and fold-change (**C**) compared to untreated ctrl and cKO mice. Non-recombined mature oligodendrocyte (MyRF^+^GFP^-^, **D**) cell density and fold-change (**E**). Data are represented as mean ± SEM and derive from N=3-4 mice (n=12 FOVs). In **B** and **D**, data from untreated groups are overlapped as dotted lines (ctrl, light green; cKO, dark green). Data were analyzed using an unpaired *t* test. For each group in **C** and **E**, data were tested using a one sample *t* test to compare their mean with the hypothetical value μ=1.

**Supplementary Figure 5.**
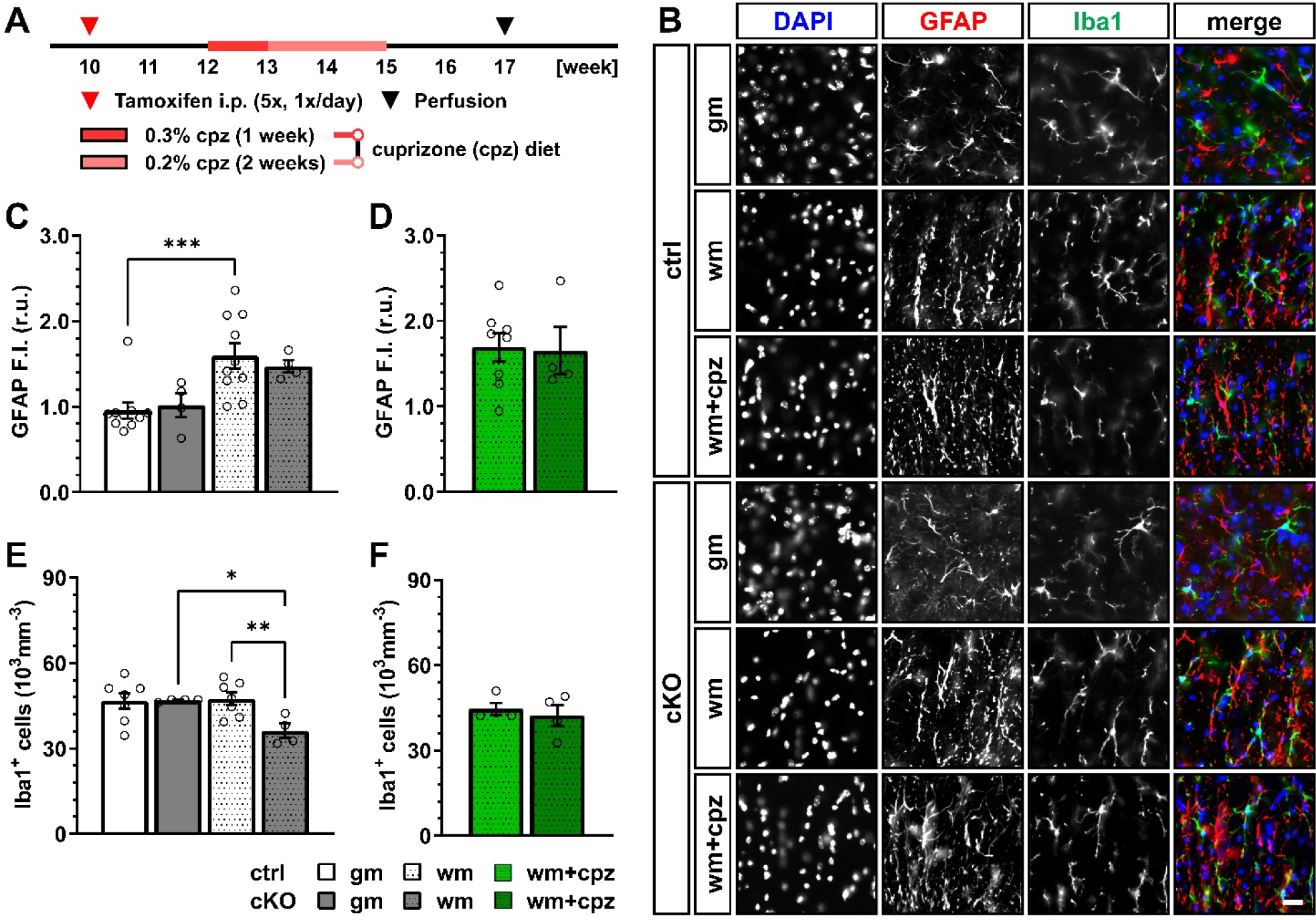
No glial response in terms of astroglial GFAP expression or microglia number in the spinal cord after cuprizone treatment. **(A)** Experimental design for tamoxifen-induced GABA_B_R deletion and GCaMP3 expression in NG2^+^ OPCs, cuprizone (cpz) treatment, perfusion and IHC analysis. (**B**) IHC of spinal cord tissue from untreated and cuprizone-treated ctrl and cKO mice (gray matter, gm; white matter, wm) stained for DAPI (blue), Glial Fibrillary Acidic Protein (GFAP, red) and Iba1 (green). Scale bar, 20 µm. Mean fluorescence intensity (F.I.) of GFAP signal in untreated (**C**) and cpz-treated (**D**) mice. Iba1^+^ cell density in untreated (**E**) and cuprizone-treated (**F**) mice. Data are represented as mean ± SEM and derive from N=3-8 mice (n=12-32 FOVs). Data were analyzed using a two-way ANOVA with multiple comparisons test (**C**, **E**) or an unpaired *t* test (**D**, **F**).

**Supplementary Figure 6.**
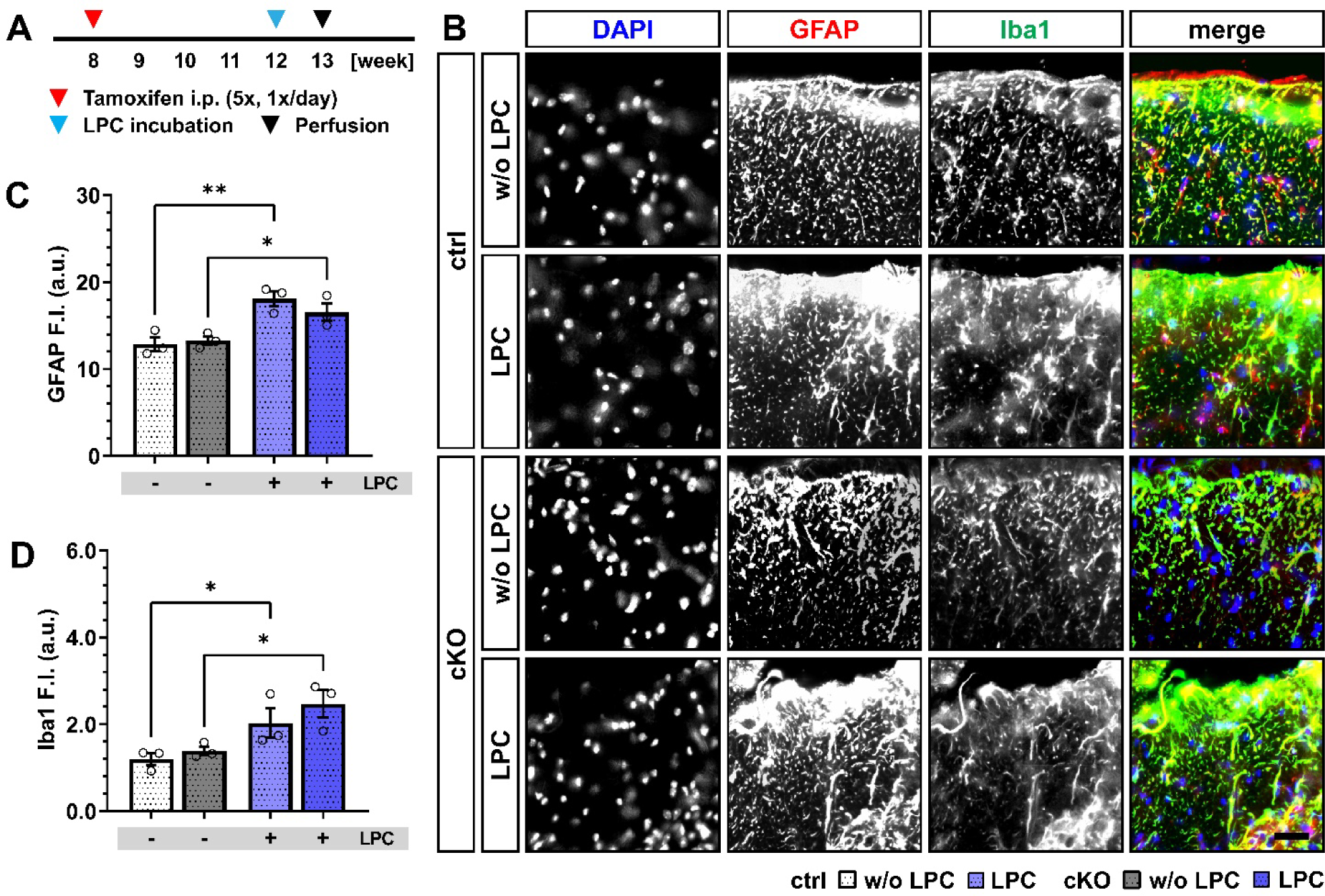
Acute LPC incubation induces strong glial response in the white matter of the spinal cord of control and cKO mice. **(A)** Experimental design for tamoxifen-induced GABA_B_R deletion and GCaMP3 (GFP) expression in NG2^+^ OPCs, lysolecithin (LPC) incubation, perfusion and IHC. (**B**) IHC of the dorsal white matter of spinal cord tissue from untreated and LPC-treated ctrl and cKO mice stained for DAPI (blue), GFAP (red) and Iba1 (green). Scale bar, 20 µm. (**C**) Mean fluorescence intensity (F.I.) of GFAP signal in LPC-treated mice. (**D**) Mean fluorescence intensity (F.I.) of Iba1 signal in LPC-treated mice. Data are represented as mean ± SEM and derive from N=3 mice (n=12 FOVs). Data were analyzed using a two-way ANOVA with multiple comparisons test.

**Supplementary Table 1.**
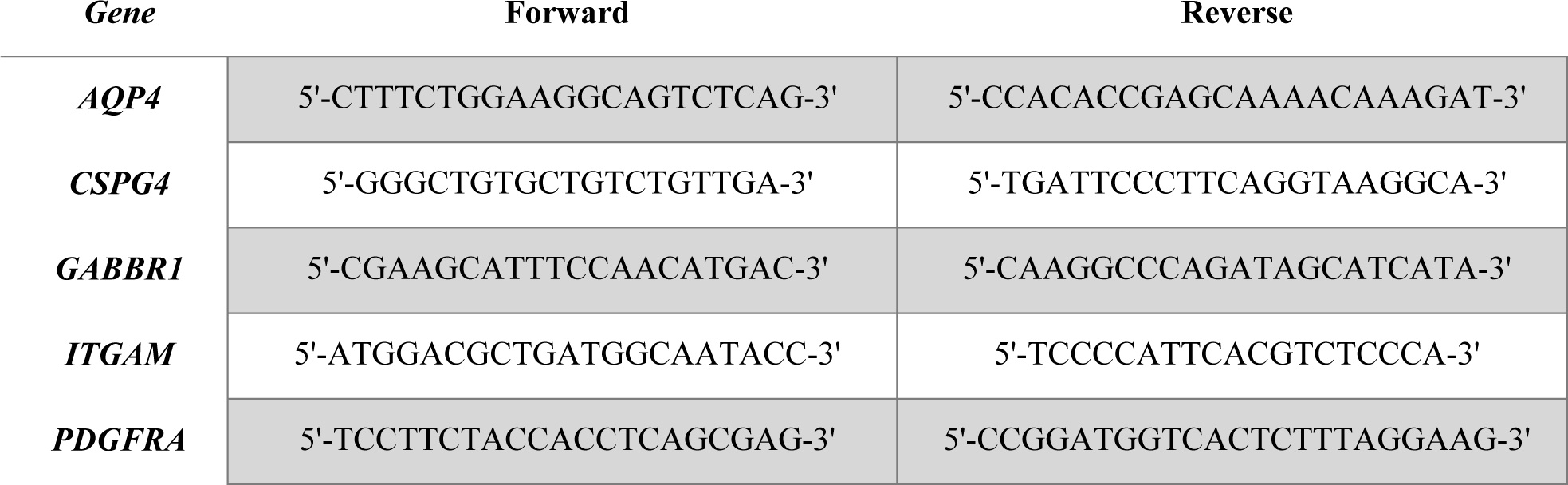
Primers used for qRT-PCR. Oligonucleotides are listed in the 5’-3’ direction.

